# Bactopia: a flexible pipeline for complete analysis of bacterial genomes

**DOI:** 10.1101/2020.02.28.969394

**Authors:** Robert A. Petit, Timothy D. Read

## Abstract

Sequencing of bacterial genomes using Illumina technology has become such a standard procedure that often data are generated faster than can be conveniently analyzed. We created a new series of pipelines called Bactopia, built using Nextflow workflow software, to provide efficient comparative genomic analyses for bacterial species or genera. Bactopia consists of a dataset setup step (Bactopia Datasets; BaDs) where a series of customizable datasets are created for the species of interest; the Bactopia Analysis Pipeline (BaAP), which performs quality control, genome assembly and several other functions based on the available datasets and outputs the processed data to a structured directory format; and a series of Bactopia Tools (BaTs) that perform specific post-processing on some or all of the processed data. BaTs include pan-genome analysis, computing average nucleotide identity between samples, extracting and profiling the 16S genes and taxonomic classification using highly conserved genes. It is expected that the number of BaTs will increase to fill specific applications in the future. As a demonstration, we performed an analysis of 1,664 public *Lactobacillus* genomes, focusing on *L. crispatus*, a species that is a common part of the human vaginal microbiome. Bactopia is an open source system that can scale from projects as small as one bacterial genome to thousands that allows for great flexibility in choosing comparison datasets and options for downstream analysis. Bactopia code can be accessed at https://www.github.com/bactopia/bactopia.

## Introduction

Sequencing a bacterial genome, an activity that once required the infrastructure of a dedicated genome center, is now a routine task that even a small laboratory can undertake. A large number of open source software tools have been created to handle various parts of the process of using raw read data for functions such as SNP calling and de novo assembly. As a result of dedicated community efforts, it has recently become much easier to locally install these bioinformatic tools through package managers (Bioconda (1), Brew) or through the use of software containers (Docker, Singularity). Despite these advances, producers of bacterial sequence data face a bewildering array of choices when considering how to perform analysis, particularly when large numbers of genomes are involved and processing efficiency and scalability become major factors.

Efficient bacterial multi-genome analysis has been hampered by three missing functionalities. First, is the need to have “*workflows of workflows*’’ that can integrate analyses and provide a simplified way to start with a collection of raw genome data, remove low quality sequences and perform the basic analytic steps of de novo assembly, mapping to reference sequence and taxonomic assignment. Second, is the desire to incorporate user-specific knowledge of the species into the input of the main genome analysis pipeline. While many microbiologists are not expert bioinformaticians, they are experts in the organisms they study. Third, is the need to create an output format from the main pipeline that could be used for future customized downstream analysis such as pan-genome analysis and basic visualization of phylogenies.

Here we introduce Bactopia, an integrated suite of workflows for flexible analysis of Illumina genome sequencing projects of bacteria from the same taxon. Bactopia is based on the Nextflow workflow software (2), and is designed to be scalable, allowing projects as small as a single genome to be run on a local desktop, or many thousands of genomes to be run as a batch on a cloud infrastructure. Running multiple tasks on a single platform standardizes the underlying data quality used for gene and variant calling between projects run in different laboratories. This structure also simplifies the user experience. In Bactopia, complex multi-genome analysis can be run in a small number of commands. However, there are myriad options for fine-tuning datasets used for analysis and the functions of the system. The underlying Nextflow structure ensures reproducibility. To illustrate the functionality of the system we performed a Bactopia analysis of 1,664 public genome projects of the *Lactobacillus* genus, an important component of the microbiome of humans and animals.

## Design and implementation

Bactopia links together open source bioinformatics software, available from Bioconda (1), using Nextflow (2). Nextflow was chosen for its flexibility: Bactopia can be run locally, on clusters, or on cloud platforms with simple parameter changes. It also manages the parallel execution of tasks and creates checkpoints allowing users to resume jobs. Nextflow automates installation of the component software of the workflow through integration with Bioconda. For ease of deployment, Bactopia can either be installed through Bioconda, a Docker container, or a Singularity container. All the software programs used by Bactopia (version v1.3.0) described in this manuscript are listed in **Table 1** with their individual version numbers.

**Table 1.**
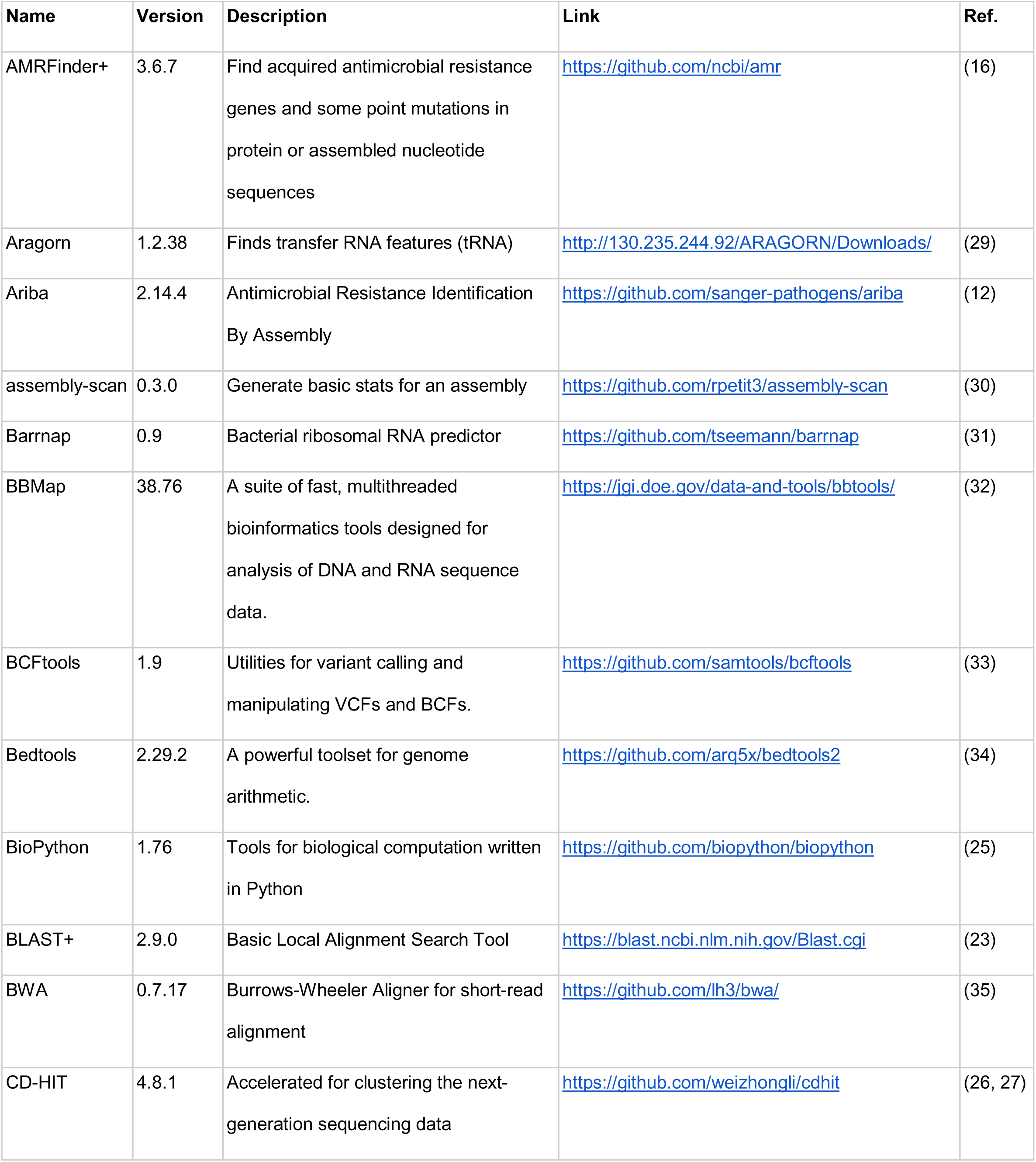
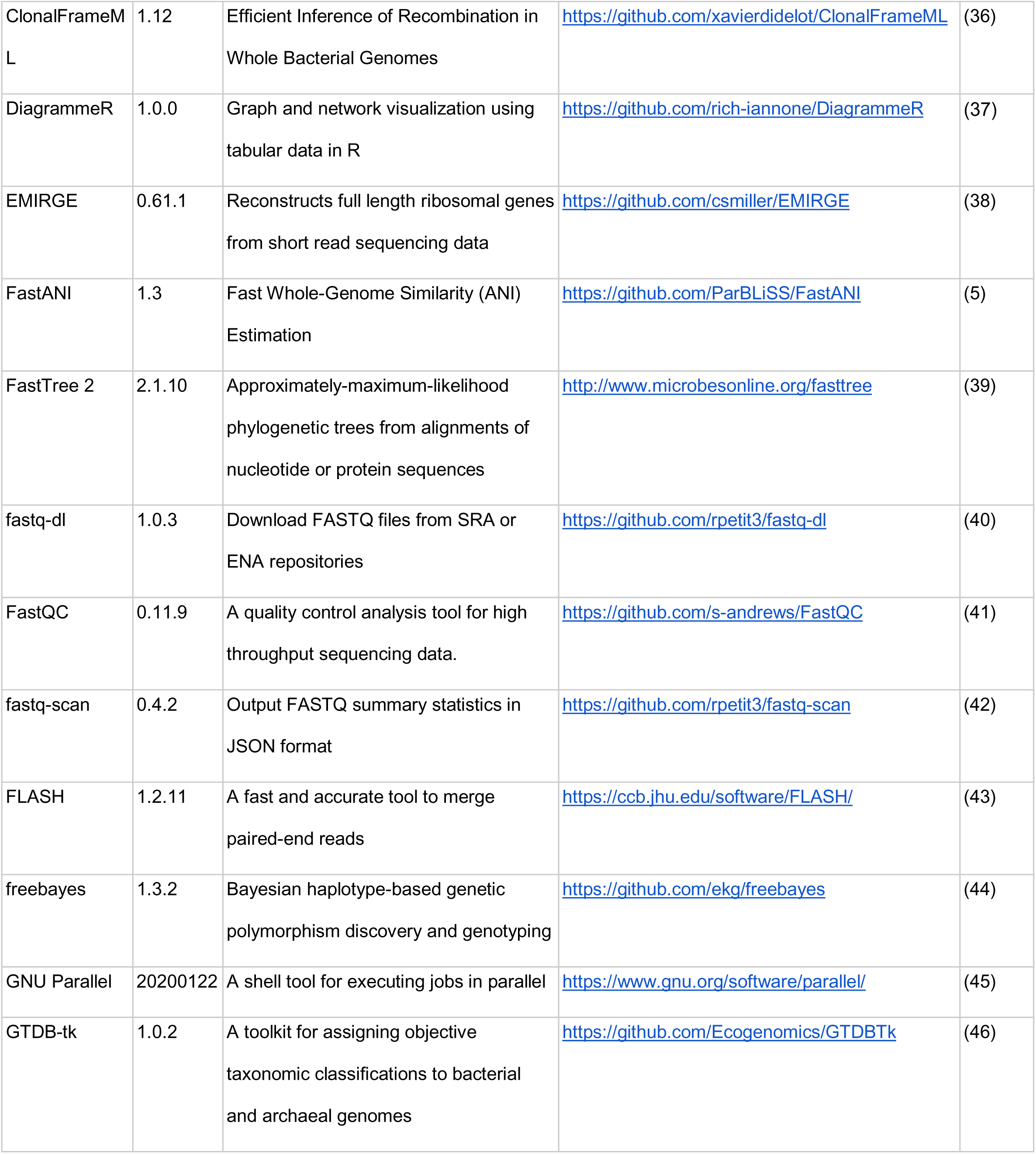
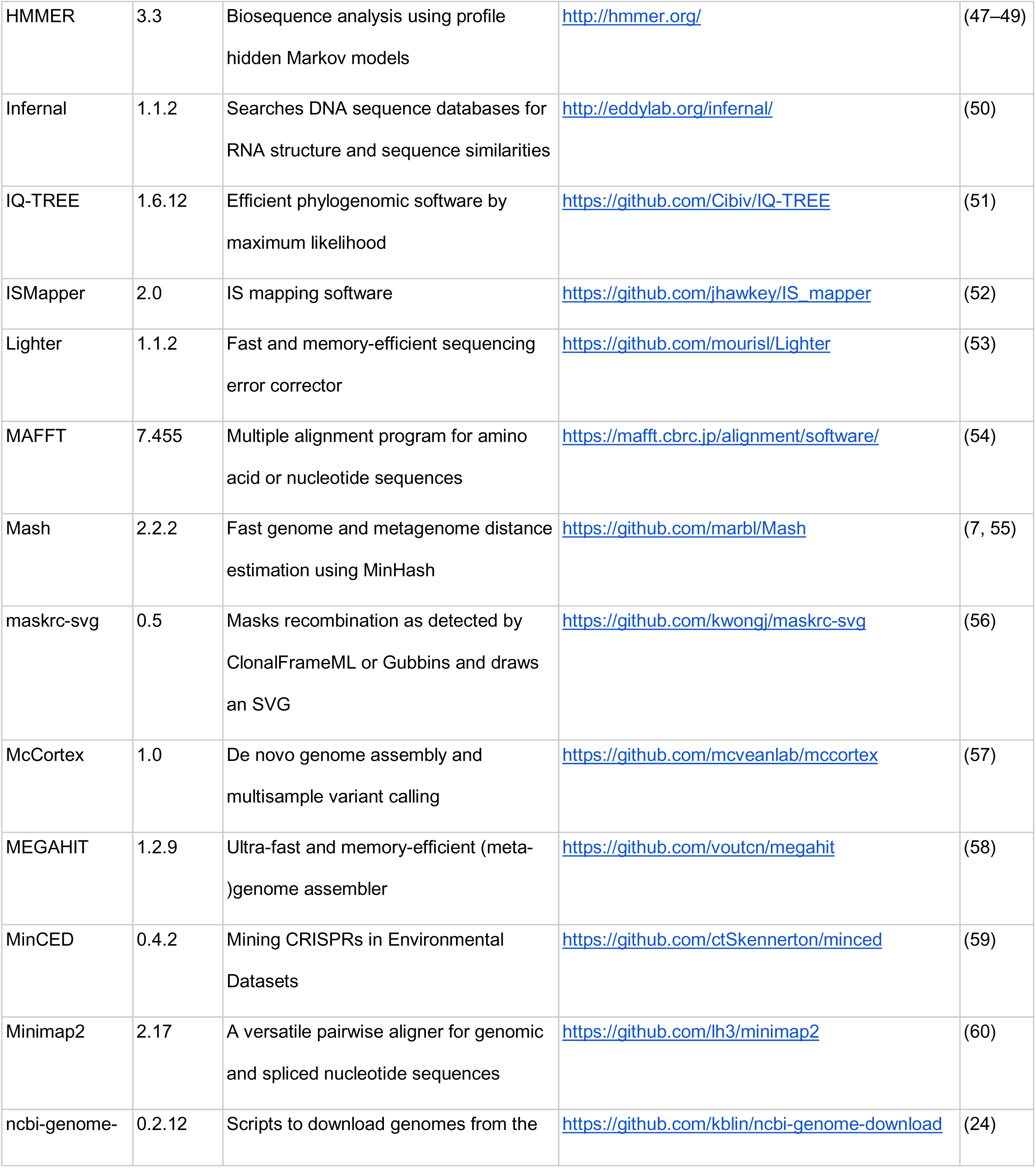
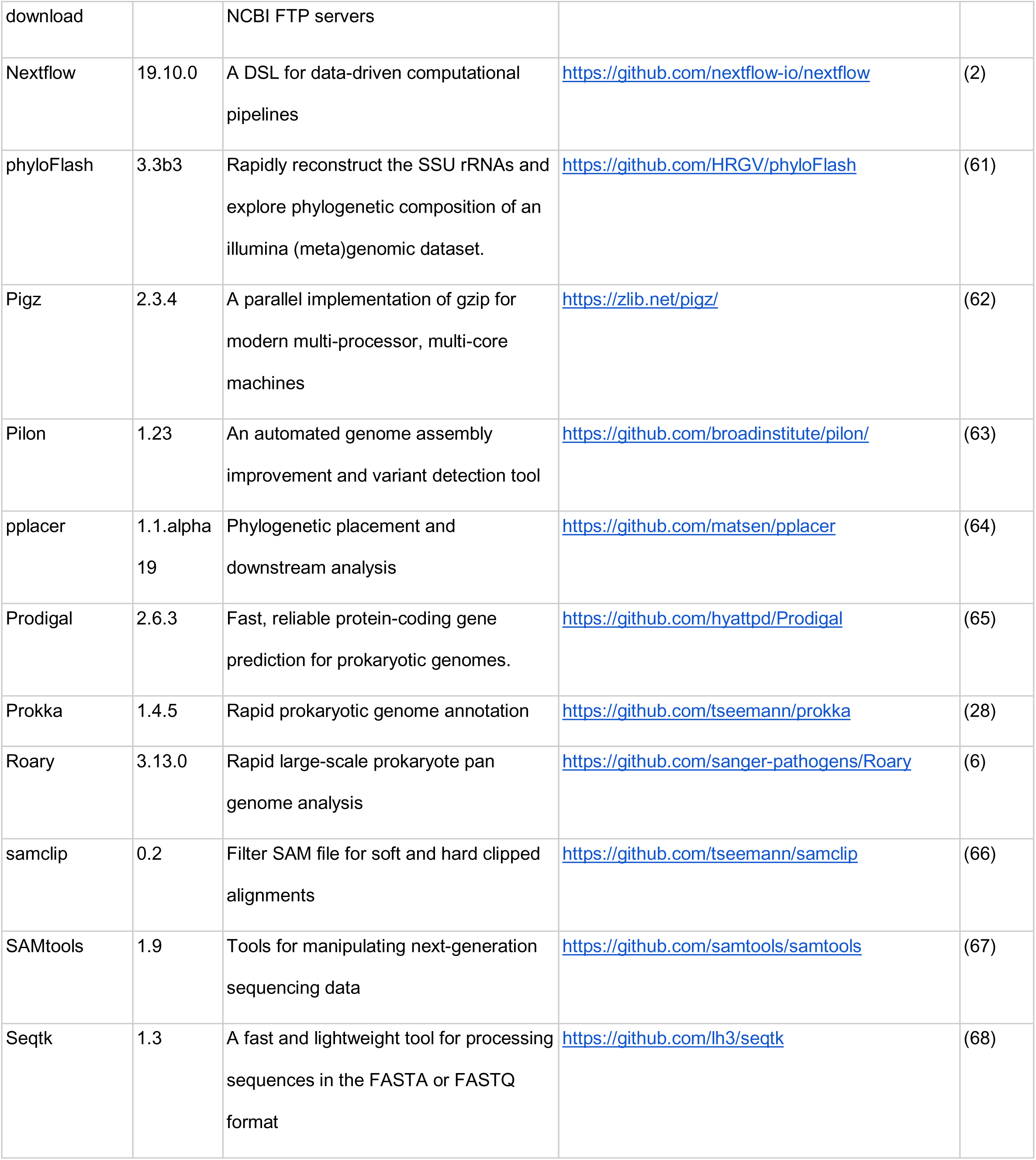
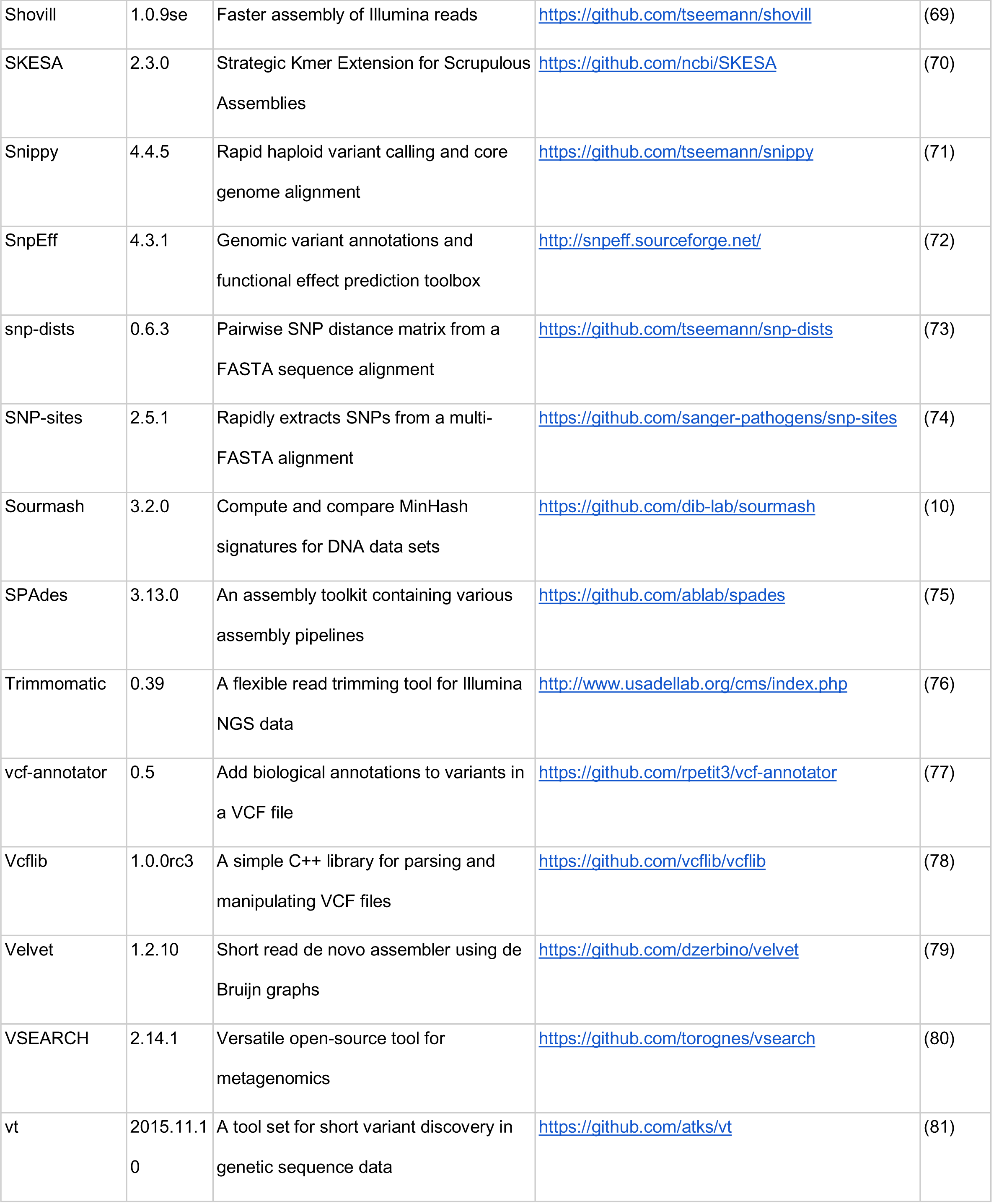
List of bioinformatic tools used by Bactopia. A list of bioinformatic tools used throughout version 1.3.0 of the Bactopia Analysis Pipeline and Bactopia Tools.

There are three main components of Bactopia (**Figure 1**): 1) <u>Bactopia Datasets (BaDs)</u>, a framework for formatting organism-specific datasets to be used by the downstream analysis pipeline; 2) <u>Bactopia Analysis Pipeline (BaAP)</u>, a customizable workflow for the analysis of individual bacterial genome projects that is an extension and generalization of the previously published *Staphylococcus aureus*-specific “Staphopia Analysis Pipeline” (StAP) (3). The inputs to BaAP are FASTQ files from bacterial Illumina sequencing projects, either imported from the National Centers for Biotechnology Information (NCBI) Short Read Archive (SRA) database or provided locally, and any reference data in the BaDs; 3) <u>Bactopia Tools (BaTs)</u>, a set of workflows that use the output files from a BaAP project to run genomic analysis on multiple genomes. For this project we used BaTs to a) summarize the results of running multiple bacterial genomes through BaAP, b) extract 16S sequences and create a phylogeny, c) assign taxonomic classifications with the Genome Taxonomy Database (4), d) subset *Lactobacillus crispatus* samples by average nucleotide identity (ANI) with FastANI (5), e) run pan-genome analysis for *L. crispatus* using Roary (6) and create a core-genome phylogeny.

**Figure 1.**
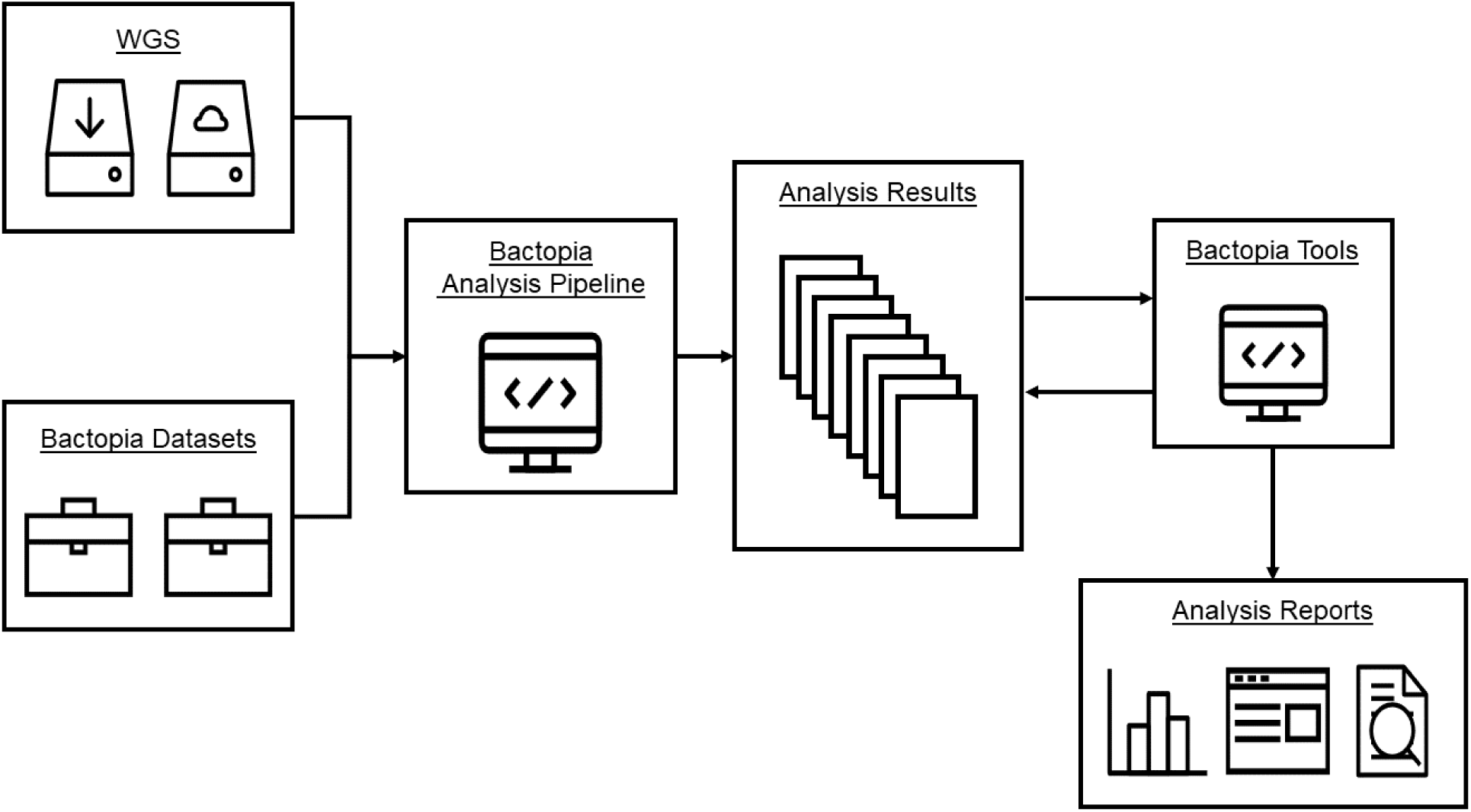
Bactopia overview.

### Bactopia Datasets

The Bactopia pipeline can be run without downloading and formatting Bactopia Datasets (BaDs). However, providing them enriches the downstream analysis. Bactopia can import specific existing public datasets, as well as accessible user-provided datasets in the appropriate format. A subcommand (*bactopia datasets*) was created to automate downloading, building, and (or) configuring these datasets for Bactopia.

BaDs can be grouped into those that are general and those that are user-supplied. General datasets include: a Mash (7) sketch of the NCBI RefSeq (8) and PLSDB (9) databases and a Sourmash (10) signature of microbial genomes (includes viral and fungal) from the NCBI GenBank (11) database. Ariba (12), a software for detecting genes in raw read (FASTQ) files, uses a number of default reference databases for virulence and antibiotic resistance. The available Ariba datasets include: ARG-ANNOT (13), CARD (14), MEGARes (15), NCBI Reference Gene Catalog (16), plasmidfinder (17), resfinder (18), SRST2 (19), VFDB (20), and VirulenceFinder (21).

When an organism name is provided, additional datasets are set up. If a MLST schema is available for the species, it is downloaded from PubMLST.org (22) and set up for BLAST+ (23) and Ariba. Each RefSeq completed genome for the species is downloaded using ncbi-genome-download (24). A Mash sketch is created from the set of downloaded completed genomes to be used for automatic reference selection for variant calling. Protein sequences are extracted from each genome with BioPython (25), clustered using CD-HIT (26, 27) and formatted to be used by Prokka (28) for annotation. Users may also provide their own organism-specific reference datasets to be used for BLAST+ alignment, short-read alignment, or variant calling.

### Bactopia Analysis Pipeline

The Bactopia Analysis Pipeline (BaAP) takes input FASTQ files and optional user-specified BaDs and performs a number of workflows that are based on either *de novo* whole genome assembly, reference mapping or sequence decomposition (ie kmer-based approaches) (**Figure 1**). BaAP has incorporated numerous existing bioinformatic tools (**Table 1**) into its workflow (**Supplementary Figure 1**). For each tool, many of the input parameters are exposed to the user, allowing for fine-tuning analysis.

### BaAP: Acquiring FASTQs

Bactopia provides multiple ways for users to provide their FASTQ formatted sequences. Input FASTQs can be local or downloaded from public repositories.

Local sequences can be processed one at a time or in batches. To process a single sample, the user provides the path to their FASTQ(s) and a sample name. For multiple samples, this method does not make efficient use of Nextflow’s queue system. Alternatively, users can provi de a “*file of filenames*” (FOFN), which is a tab-delimited file with information about samples and paths to the corresponding FASTQ(s). By using the FOFN method, Nextflow queues each sample and makes efficient use of available resources. A subcommand (*bactopia prepare*) was created to automate the creation of a FOFN.

Raw sequences available from public repositories (e.g. European Nucleotide Archive (ENA), Sequence Read Archive (SRA), or DNA Data Bank of Japan (DDBJ)) can also be processed by Bactopia. Sequences associated with a provided Experiment accession (e.g. DRX, ERX, SRX) are downloaded and processed exactly as local sequences would be. A subcommand (*bactopia search*) was created, which allows users to query ENA to create a list of Experiment accessions from the ENA Data Warehouse API (82) associated with a BioProject accession, Taxon ID, or organism name.

### BaAP: Validating FASTQs

The path for input FASTQ(s) is validated, and if necessary, sequences from public repositories are downloaded using fastq-dl (40). Once validated, the FASTQ input(s) are tested to determine if they meet a minimum threshold for continued processing. All BaAP steps expect to use Illumina sequence data, which is the great majority of genome projects currently generated. FASTQ files that are explicitly marked as non-Illumina or have properties that suggest non-Illumina (e.g read-length, error profile) are excluded. By default, input FASTQs must exceed 2,241,820 bases (20x coverage of the smallest bacterial genome, *Nasuia deltocephalinicola* (83)) and 7,472 reads (minimum required base pairs / 300 bp, the longest available reads from Illumina). If estimated, the genome size must be between 100,000 bp and 18,040,666 bp which is based off the range of known bacterial genome sizes (*N. deltocephalinicola,* GCF_000442605, 112,091 bp*; Minicystis rosea*, GCF_001931535, 16,040,666 bp). Failure to pass these requirements excludes the samples from further subsequent analysis. The threshold values can be adjusted by the user at runtime.

### BaAP: FastQ Quality Control and generation of pFASTQ

Input FASTQs that pass the validation steps undergo quality control steps to remove poor quality reads. BBDuk, a component of BBTools (32), removes Illumina adapters, phiX contaminants and filters reads based on length and quality. Base calls are corrected using Lighter (53). At this stage, the default procedure is to downsample the FASTQ file to an average 100x genome coverage (if over 100x) with Reformat (from BBTools). This step, which was used in StAP (3), significantly saves compute time at little final cost to assembly or SNP calling accuracy. The genome size for coverage calculation is either provided by the user or estimated based on the FASTQ data by Mash (7). The user can provide their own value for downsampling FASTQs or disable it completely. Summary statistics before and after QC are created using FastQC (41) and fastq-scan (42). After QC, the original FASTQs are no longer used and only the processed FASTQs (pFASTQ) are used in subsequent analysis.

### BaAP: Assembly, Reference Mapping and Decomposition

BaAP uses Shovill (69) to create a draft *de novo* assembly with MEGAHIT (58), SKESA (70) (default), SPAdes (75) or Velvet (79) and make corrections using Pilon (63) from the pFASTQ. Summary statistics for the draft assembly are created using assembly-scan (30). If the total size of the draft assembly fails to meet a user-specified minimum size, further assembly-based analyses are discontinued. Otherwise, a BLAST+ (23) nucleotide database is created from the contigs. The draft assembly is also annotated using Prokka (28). If available at runtime, Prokka will first annotate with a clustered RefSeq protein set, followed by its default databases. The annotated genes and proteins are then subjected to antimicrobial resistance prediction with AMRFinder+ (16).

For each pFASTQ, sketches are created using Mash (*k*=21,31) and Sourmash (10) (*k*=21,31,51). McCortex (57) is used to count 31-mers in the pFASTQ.

### BaAP: Optional Steps

At runtime, Bactopia checks for BaDs specified by the command line (if any) and adjusts the settings of the pipeline accordingly. Examples of processes executed only if a BaDs is specified include Ariba (12) analysis for each available reference dataset; sequence containment screening against RefSeq (8) with mash screen (55) and against GenBank (11) with sourmash lca gather (10), and PLSDB (9), with mash screen and BLAST+. The sequence type (ST) of the sample is determined with BLAST+ and Ariba. The nearest reference RefSeq genome, based on mash (7) distance, is downloaded with ncbi-genome-download (24), and variants are called with Snippy (71). Alternatively, one or more reference genomes can be provided by the user. Users can also provide sequences for: sequence alignment and per-base coverage with BWA (35, 84) and Bedtools (34), BLAST+ alignment, or insertion site determination with ISMapper (52).

### Bactopia Tools

After BaAP has successfully completed, it will create a directory for each strain with subdirectories for each analysis result. The directory structure is independent of project or options chosen. Bactopia Tools (BaTs) are a set of comparative analysis workflows written using Nextflow that take advantage of the predictable output structure from BaAP. Each BaT is created from the same framework and a subcommand (*bactopia tools create*) is available to simplify the creation of future BaTs.

Five BaTs were used for analyses in this article. The *summary* BaT, outputs a summary report of the set of samples and a list of samples that failed to meet thresholds set by the user. This summary includes basic sequence and assembly stats as well as technical (pass/fail) information. The *roary* BaT creates a pan-genome of the set of samples with Roary (6), with the option to include RefSeq (8) completed genomes. The *fastani* BaT determines the pairwise average nucleotide identity (ANI) for each sample with FastANI (5). The *phyloflash* BaT reconstructs 16S rRNA gene sequences with phyloFlash (61). The *gtdb* BaT assigns taxonomic classifications from the Genome Taxonomy Database (GTDB) (4) with GTDB-tk (46). Each Bactopia tool has a separate Nextflow workflow with its own conda environment, Docker image and Singularity image.

## Comparison to Similar Open Source Software

At the time of writing (February 2020), we knew of only three other actively-maintained open source generalist bacterial genomic workflow softwares that encompassed a similar range of functionality to Bactopia: ASA^3^P (85), TORMES (86) and the currently unpublished Nullarbor (87). The versions of these programs used many of the same component software (e.g. Prokka, SPAdes, BLAST+, Roary) but differed in the philosophies underlying their design (**Table 2**). This made head-to-head runtime comparisons somewhat meaningless, as each was aimed at different analysis scenarios and produced different output. Bactopia was the most open-ended and flexible, allowing the user to customize input databases and providing a platform for downstream analysis by different BaTs rather than built-in pangenome and phylogeny creation. Bactopia also had some features not implemented in the others, such as SRA/ENA search and download and automated reference genome selection for identifying variants. Both Bactopia and ASA^3^P are highly scalable, each can be seamlessly executed on local, cluster, and cloud environments with little effort required by the user. ASA^3^P was the only program to implement long-read data assembly. TORMES was the only program to include a user customizable RMarkdown for report and to have optional analyses specifically for *Escherichia* and *Salmonella*. Nullarbor was the only program to implement a pre-screen method for filtering out potential biological outliers prior to full analysis.

**Table 2.** A comparison of bacterial genome analysis workflows.

| Feature | Bactopia | ASA<sup>3</sup>P | Nullarbor | TORMES |
| --- | --- | --- | --- | --- |
| Version | 1.3.0 | 1.2.2 | 2.0.20191013 | 1.0 |
| Release Date | February 19th, 2020 | November 20th, 2019 | October 13th, 2019 | February 4th, 2019 |
| Latest Commit | February 23th, 2020 | February 19th, 2020 | February 23th, 2020 | January 30th, 2020 |
| Sequence Technology | Illumina | Illumina, Nanopore, PacBio | Illumina | Illumina |
| Single End Reads | Yes | Yes | No | No |
| Workflow | Nextflow | Groovy | Perl + Make | Bash |
| Resume If Stopped | Yes | No | Yes | No |
| Reuse Existing Runs For Expanded Analysis | Yes | No | Yes | No |
| Built In HPC Cluster and Cloud Capability | Yes | Yes | No | No |
| Individual Program Adjustable Parameters | Yes | No | Yes | No |
| Batch processing from Config File | Yes | Yes | Yes | Yes |
| Single Sample Processing from Command Line | Yes | No | Yes | No |
| Sequence Depth<br>Downsample | Yes | No | Yes | No |
| Automatic<br>Reference<br>Selection For<br>Variant Detection | Yes | No | No | No |
| Data Download<br>from SRA/ENA | Yes | No | No | No |
| Species<br>Identification | *k*-mers,16S, ANI | *k*-mers, 16S, ANI | *k*-mers | *k*-mers |
| Comparative<br>Analysis | Separate Process | Built In Process | Built In Process | Built In Process |
| Summary | Text | HTML | HTML | R Markdown |
| Package Manager | Bioconda | - | Bioconda & Brew | Conda YAML |
| Container Available | Yes | Yes | Yes | No |
| Documentation | Website | PDF Manual | README | README |
| Github Repository | [bactopia/bactopia](https://github.com/bactopia/bactopia) | [oschwengers/asap](https://github.com/oschwengers/asap) | [tseemann/nullarbor](https://github.com/tseemann/nullarbor) | [nmquijada/tormes](https://github.com/nmqijada/tormes) |

## Use case: the *Lactobacillus* genus

We performed a Bactopia analysis of publicly-available raw Illumina data labelled as belonging to the *Lactobacillus* genus. *Lactobacillus* is an important component of the human microbiome, and cultured samples have been sequenced by several research groups over the past few years. *L. crispatus* and other species are often the majority bacterial genus of the human vagina and are associated with low pH and reduction in pathogen burden (88). Samples of the genus are used in the food industry for fermentation in the production of yoghurt, kimchi, kombucha and other common items. *Lactobacillus* is a common probiotic, although recent genomic-based transmission studies showed that bloodstream infections can follow after ingestion by immunocompromised patients (89).

### Running BaAP

In November 2019, we initiated Bactopia analysis using the following three commands:

~~~
# Build Lactobacillus dataset
bactopia datasets ∼/bactopia-datasets --species ‘Lactobacillus’ –
include_genus --cpus 10

# Query ENA for all Lactobacillus (tax id 1578) sequence projects
bactopia search 1578 –prefix lactobacillus
# this creates a file called ‘lactobacillus-accessions.txt’

# Process Lactobacillus samples
mkdir ∼/bactopia
cd ∼/bactopia
bactopia --accessions ∼/lactobacillus-accessions.txt --datasets ∼/bactopia-
datasets --species lactobacillus --coverage 100 --cpus 4 --min_genome_size
1000000 --max_genome_size 4200000
~~~

The *bactopia datasets* subcommand automated downloading of BaDs. With these parameters, we downloaded and formatted the following datasets: Ariba (12) reference databases for the Comprehensive Antibiotic Resistance Database (CARD) and the core Virulence Factor Database (VFDB) (14, 20), RefSeq Mash sketch (7, 8), GenBank Sourmash signatures (10, 11), PLSDB blast database and sketch (9), and a clustered protein set and Mash sketch from completed Lactobacillus genomes available from NCBI Assembly (RefSeq). This took 25 minutes to complete.

The *bactopia search* subcommand produced a list of 2,030 Experiment accessions that had been labelled as “Lactobacillus” (taxonomy ID: 1578) (**Supplementary Data 1**). After filtering for only Illumina sequencing, 1,664 Experiment accessions remained (**Supplementary Data 2**).

The main *bactopia* command automated BaAP processing of the list of accessions using the downloaded BaTs. Here we chose a standard maximum coverage per genome of 100x, based on the estimated genome size. We used the range of genome sizes (1.2 Mb to 3.7Mb) for the completed Lactobacillus genomes to require that the estimated genome size for each sample be between 1Mbp and 4.2 Mbp.

Samples were processed on a 96-core SLURM cluster with 512 GB of available RAM. Analysis took approximately 2.5 days to complete, with an estimated runtime of 30 minutes per sample, (determined by adding up the median process runtime, 17 total, in BaAP). No individual process used more than 8GB of memory, with all but 5 using less than 1 GB. Nextflow (2) recorded detailed statistics on resource usage, including CPU, memory, job duration and I/O. (**Supplementary Data 3**).

### Analysis of *Lactobacillus* genomes using BaTs

The BaAP outputted a directory of directories named after the unique Experiment accessions for each sample. Within each sample directory were subdirectories for the output of each analysis run. This data structure was recognized by BaTs for subsequent analysis.

We used BaT *summary* to generate a summary report of our analysis. The report included an overview of sequence quality, assembly statistics, and predicted antimicrobial resistances and virulence factors. It also output a list of samples that failed to meet minimum sequencing depth and/or quality thresholds.

~~~
bactopia tools summary --bactopia ∼/bactopia --prefix lactobacillus
~~~

BaT *summary* grouped samples as “Gold”, “Silver”, “Bronze”, “Exclude” or “QC Failure”, based on BaAP completion, minimum sequencing coverage, per-read sequencing mean quality, minimum mean read length, and assembly quality (**Table 3; Supplementary Figure 2**). We used the default values for these cutoffs to group our samples. Gold samples had greater than 100x coverage, per-read mean quality greater than Q30, mean read length greater than 95bp, and an assembly with less than 100 contigs. Silver samples had greater than 50x coverage, per-read mean quality greater than Q20, mean read length greater than 75bp, and an assembly with less than 200 contigs. Bronze samples had greater than 20x coverage, per-read mean quality greater than Q12, mean read length greater than 49bp, and an assembly with less than 500 contigs. 106 samples (Exclude and QC Failure) were excluded from further analysis (**Supplementary Table 1**). 48 samples that failed to meet the minimum thresholds for Bronze quality were grouped as Exclude. 58 samples that were not processed by BaAP due to sequencing related errors or the estimated genome size were grouped as QC Failure. Of these, one (Experiment accession SRX4526092) was labeled as paired-end but did not have both sets of reads, one (Experiment accession SRX1490246) was identified to be an assembly converted to FASTQ format, and 14 had insufficient sequencing depth. The remaining 42 samples, unprocessed by BaAP, had an estimated genome size which exceeded 4.2Mbp (set at runtime). We queried these samples against available GenBank and RefSeq sketches using Mash screen and Sourmash lca gather. There were 36 samples that contained evidence for *Lactobacillus* but also sequences for other bacterial species, phage, viruses and plants genomes. There were 6 samples that contained no evidence for Lactobacillus, 4 had matches to multiple bacterial species and 2 to only *Saccharomyces cerevisiae*.

**Table 3.**
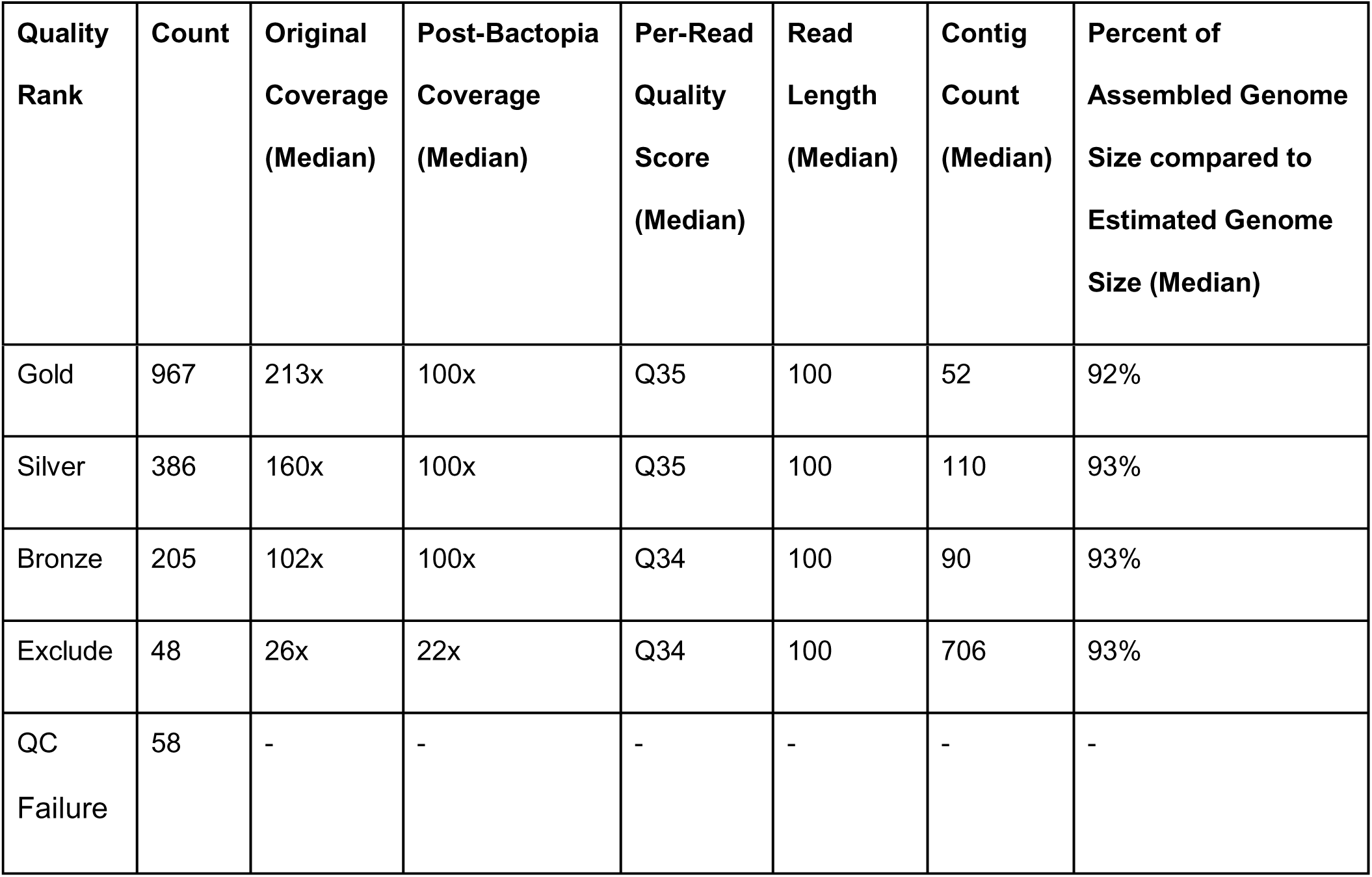
Summary of *Lactobacillus* genome sequencing projects quality and coverage.

There were 1,558 Gold, Silver or Bronze quality samples (**Table 3**) used for further analysis. For these we found that, on average, the assembled genome size was about 12% smaller than the estimated genome size (**Supplementary Figure 3**, **Table 3**). If we assume the assembled genome size is a better indicator of a sample’s genome size, the average coverage before QC increases from 220x to 268x. In this use case, the *Lactobacillus* genus, it was necessary to estimate genome sizes, but when dealing with samples from a single species it may be better to provide a known genome size.

For visualization of the phylogenetic relationships of the samples, we used the *phyloflash* and *gtdb* BaTs.

~~~
bactopia tools phyloflash --phyloflash ∼/bactopia-datasets/16s/138 --bactopia
∼/bactopia --cpus 16 --exclude ∼/bactopia-tool/summary/lactobacillus-
exclude.txt

bactopia tools gtdb --gtdb ∼/bactopia-datasets/gtdb/db --bactopia ∼/bactopia
--cpus 48 --exclude ∼/bactopia-tool/summary/lactobacillus-exclude.txt
~~~

The *gtdb* BaT used GTDB-Tk (46) to assign a taxonomic classification to each sample. GTDB-Tk used the assembly to predict genes with Prodigal (65), identify marker genes (4) for phylogenetic inference with HMMER3 (47), and find the maximum-likelihood placement of each sample on the GTDB-Tk reference tree with pplacer (64). A taxonomic classification was assigned to 1,554 samples, and 4 samples failed classification due to insufficient marker gene coverage or marker genes with multiple hits.

The *phyloflash* BaT used the phyloFlash tool (61) to reconstruct a 16S rRNA gene from each sample that was used for phylogenetic reconstruction (**Figure 2**). The 16S rRNA was reconstructed from a SPAdes (75) assembly and annotated against the SILVA (92) rRNA database (v138) database for 1,470 samples. There were 88 samples that were excluded from the phylogeny: 12 samples that did not meet the requirement of a mean read length of 50bp, 17 samples in which a 16S could not be reconstructed, 19 samples that had a mismatch in assembly and mapped read taxon designations, and 40 samples that had 16S genes reconstructed for multiple species. A phylogenetic tree was created with IQ-TREE (51, 90, 91) based on a multiple sequence alignment of the reconstructed 16S genes with MAFFT (54). Taxonomic classifications from GTDB-Tk were used to annotate the 16S with iTOL (93).

**Figure 2.**
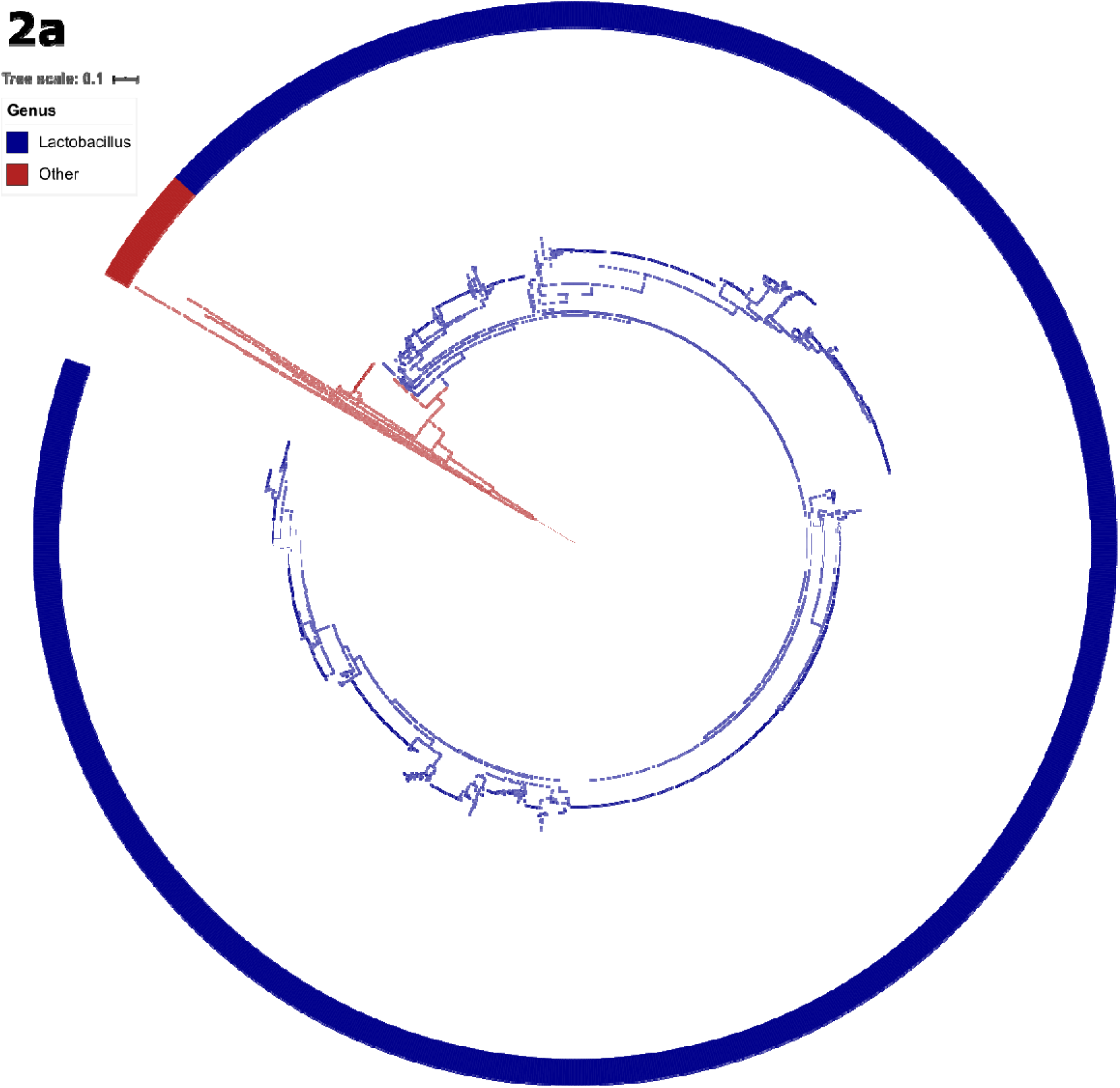
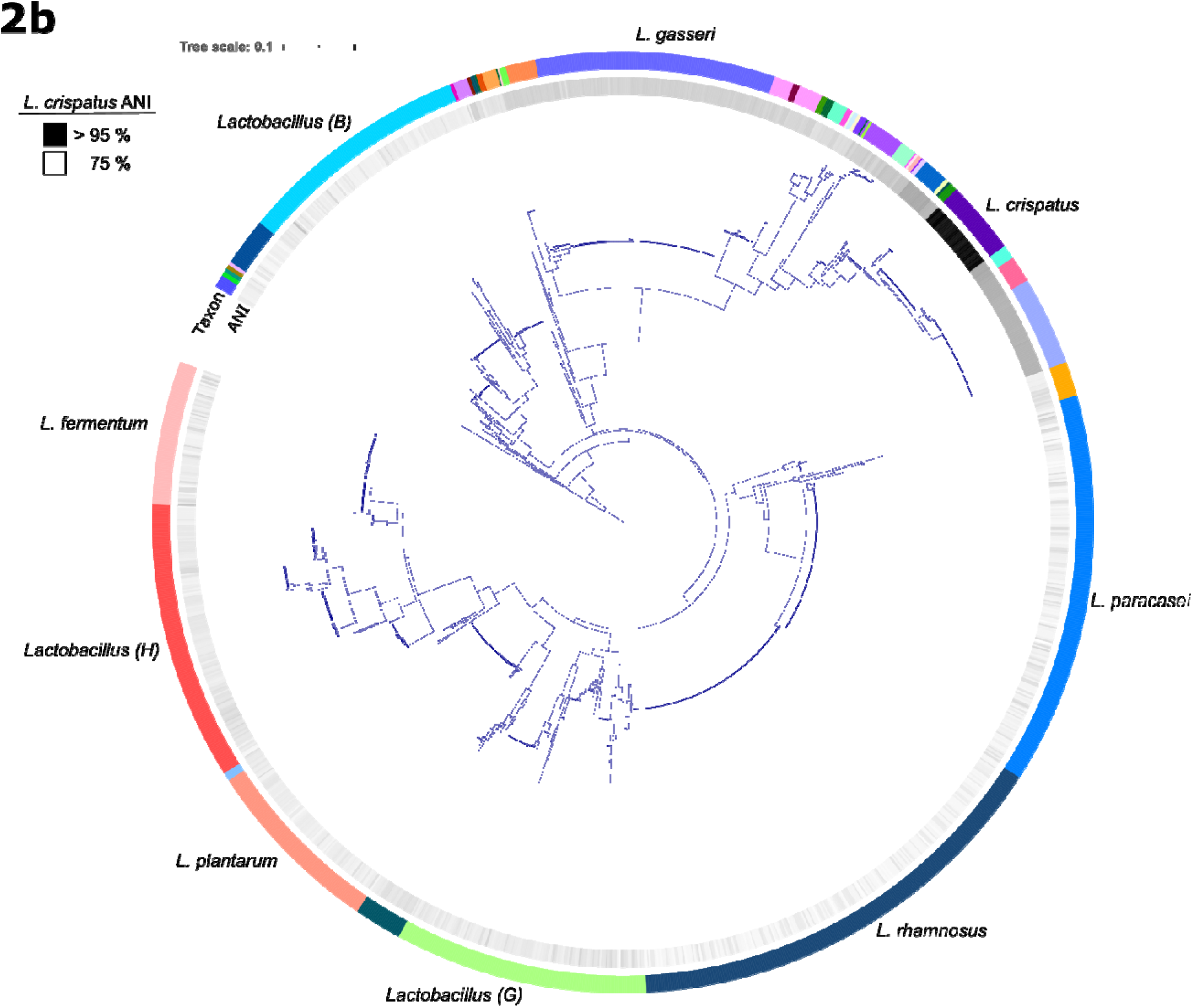
Maximum-likelihood phylogeny from reconstructed 16S rRNA genes. A phylogenetic representation 1,470 samples using IQ-Tree (51, 90, 91). A) A tree of the full set of samples. The outer ring represents the genus assigned by GTDB-Tk, with Blue being Lactobacillus and all other assignments Red. B) The same tree as A, but with the non-Lactobacillus clade collapsed. Major groups of Lactobacillus groups (depicted with a letter) and the most sequenced Lactobacillus species have been labeled. The inner ring represents the average nucleotide identity (ANI), determined by FastANI (5) of samples to *L. crispatus*. The tree was built from a multiple sequence alignment (54) of 16S genes reconstructed by phyloFlash (61) with 1,281 parsimony informative sites. The likelihood score for the consensus tree constructed from 1,000 bootstrap trees was −54,698. Taxonomic classifications were assigned by GTDB-Tk (46).

A recent analysis of completed genomes in the NCBI found 239 discontinuous *de novo Lactobacillus* species using a 94% ANI cutoff (94). Based on GTDB taxonomic classification, which applies a 95% ANI cutoff, we identified 161 distinct Lactobacillus species in 1,554 samples. The five most sequenced *Lactobacillus* species, 45% of the total, were *L. rhamnosus* (n=225), *L. paracasei* (n=180), *L. gasseri* (n=132), *L. plantarum* (n=86) and *L. fermentum* (n=80). Within these five species the assembled genomes sizes were remarkably consistent (**Supplementary Figure 4**). There were 58 samples that were not classified as Lactobacillus, of which 34 were classified as *Streptococcus pneumoniae* by both 16S and GTDB (**Supplementary Table 2**).

We found that in 505 (∼33%) out of 1,554 taxonomically classifications by 16S and GTDB were in conflict with the taxonomy according to the NCBI SRA, illustrating the importance of the unbiased approach to understanding sample context. In samples that had both a 16S and GTDB taxonomic classification, there was disagreement in 154 out of 1,467 samples. Of these, 47% were accounted for by the recently described *L. paragasseri* (95) (n=72). This possibly highlights a lag in the reclassification of assemblies in the NCBI Assembly database.

Analysis of the pangenome of the entire genus using a tool such as Roary (6), would return only a few core genes, owing to sequence divergence of evolutionary distant species. However, because the *roary* BaT can be supplied a list of individual samples it is possible to isolate the analysis to the species level. As an example of using BaTs to focus on a particular group within the larger set of results, we chose *L. crispatus*, a species commonly isolated from the human vagina and also found in the guts/feces of poultry. We used the *fastani* BaT to estimate the ANI of all samples against a randomly selected *L. crispatus* completed genome (NCBI Assembly Accession: GCF_003795065) with FastANI (5). A cutoff of greater than 95% ANI was used to categorize a sample as *L. crispatus*. A pan-genome analysis was conducted on only the samples categorized as *L. crispatus* using the *roary* BaT. The *roary* BaT downloaded all available completed *L. crispatus* genomes with ncbi-genome-download (24), formatted the completed genomes with Prokka (28), created a pan-genome with Roary (6), identified and masked recombination with ClonalFrameML (36) and maskrc-svg (56), created a phylogenetic tree with IQ-TREE (51, 90, 91) and a pair-wise SNP distance matrix with snp-dists (73).

~~~
bactopia tools fastani --bactopia ∼/bactopia --exclude ∼/bactopia-
tool/summary/lactobacillus-exclude.txt --accession GCF_003795065.1 –
refseq_only --minFraction 0.0

# Identify samples with > 95% ANI to L. crispatus
awk ‘{if ($3 > 95){print $0}}’ ∼/bactopia-tool/fastani/fastani.tsv | grep
“RX” > ∼/crispatus-include.txt

bactopia tools roary --bactopia ∼/bactopia --cpus 20 --include ∼/crispatus-
include.txt --species “lactobacillus crispatus” --n
~~~

ANI analysis revealed 38 samples as having >96.1% ANI to *L. crispatus*, with no other sample greater than 83.1%. Four completed *L. crispatus* genomes were also included in the analysis (**Table 4**) for a total of 42 genomes. The pan-genome of *L. crispatus* was revealed to have 7,037 gene families and 972 core genes (**Figure 3**). Similar to a recent analysis by Pan et al (96), *L. crispatus* was separated into two main phylogenetic groups, one associated with human vaginal isolates, the other of more mixed provenance, and including chicken, turkey and human gut isolates.

**Figure 3.**
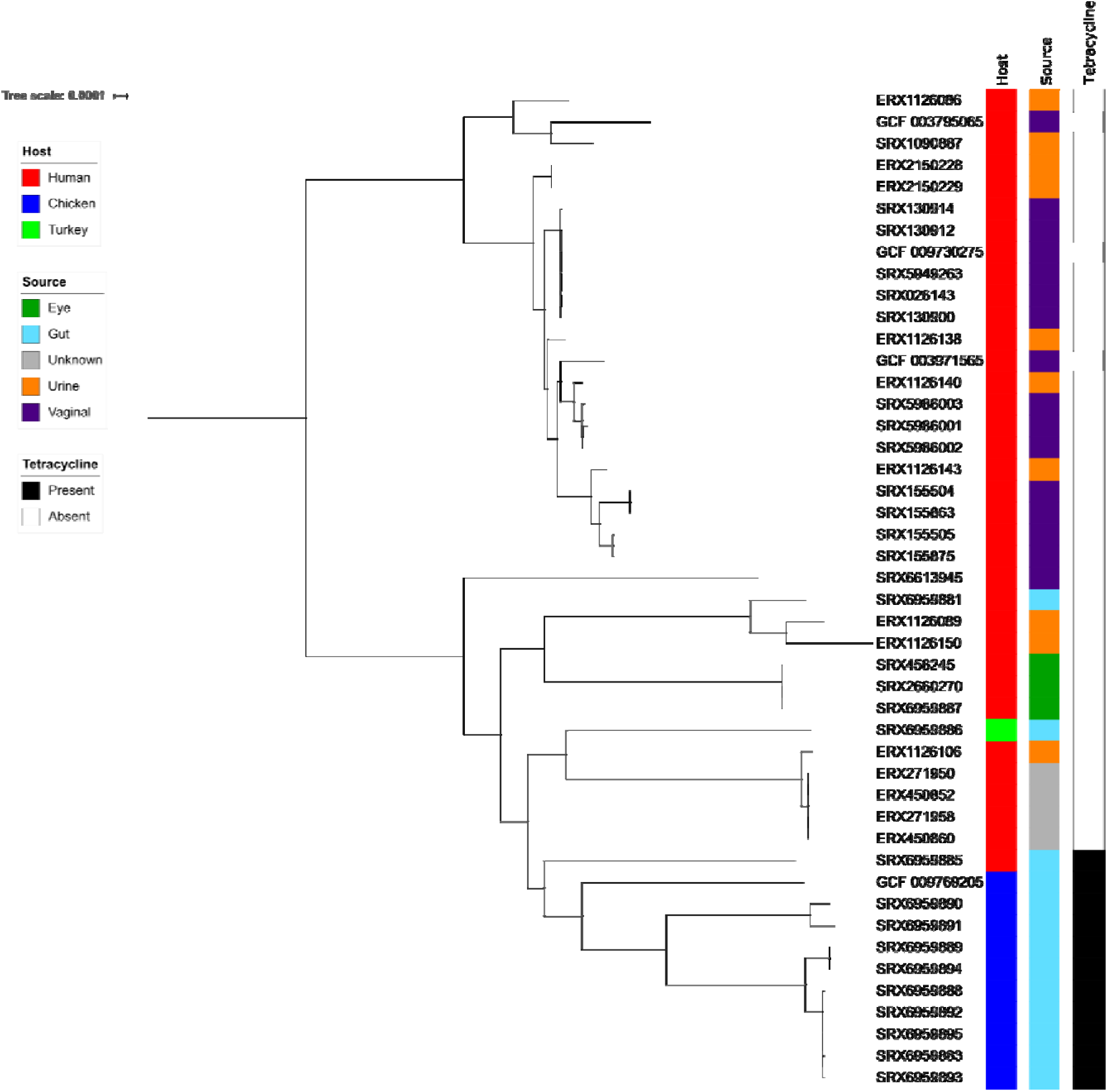
Core-genome maximum-likelihood phylogeny of *Lactobacillus crispatus*. A core-genome phylogenetic representation using IQ-Tree (51, 90, 91) of 42 *L. crispatus* samples. The putatively recombinant positions predicted using ClonalFrameML (36) were removed from the alignment with maskrc-svg (56). The tree was built from 972 core genes identified by Roary with 9,209 parsimony informative sites. The log likelihood score for the consensus tree constructed from 1,000 bootstrap trees was −1,418,106.

**Table 4.** *L crispatus* genomes used in pan-genome analysis. *Lactobacillus crispatus* samples (n=42) used in the pan-genome analysis. An accession corresponding to the BioProject, BioSample, and either the NCBI Assembly accession (beginning with GCF) or SRA Experiment accession is given for each sample. The host and source were collected from metadata associated with the BioSample or available publications. In cases when a host and/or source was not explicitly stated, it was inferred from available metadata (denoted by *).

| BioProject | BioSample | Accession | Host | Source | Reference |
| --- | --- | --- | --- | --- | --- |
| PRJEB8104 | SAMEA3319334 | ERX1126086 | Human* | urine* |  |
| PRJEB8104 | SAMEA3319350 | ERX1126089 | Human* | urine* |  |
| PRJEB8104 | SAMEA3319265 | ERX1126106 | Human* | urine* |  |
| PRJEB8104 | SAMEA3319366 | ERX1126138 | Human* | urine* |  |
| PRJEB8104 | SAMEA3319373 | ERX1126140 | Human* | urine* |  |
| PRJEB8104 | SAMEA3319383 | ERX1126143 | Human* | urine* |  |
| PRJEB8104 | SAMEA3319392 | ERX1126150 | Human* | urine* |  |
| PRJEB22112 | SAMEA104208649 | ERX2150228 | Human* | urine* |  |
| PRJEB22112 | SAMEA104208650 | ERX2150229 | Human* | urine* |  |
| PRJEB3060 | SAMEA1920319 | ERX271950 | Human* | unknown |  |
| PRJEB3060 | SAMEA1920326 | ERX271958 | Human* | unknown |  |
| PRJEB3060 | SAMEA1920319 | ERX450852 | Human* | unknown |  |
| PRJEB3060 | SAMEA1920326 | ERX450860 | Human* | unknown |  |
| PRJNA50051 | SAMN00109860 | SRX026143 | Human* | vaginal* |  |
| PRJNA272101 | SAMN03854351 | SRX1090887 | Human | urine | (97) |
| PRJNA50053 | SAMN00829399 | SRX130900 | Human* | vaginal* |  |
| PRJNA50057 | SAMN00829123 | SRX130912 | Human* | vaginal* |  |
| PRJNA50067 | SAMN00829125 | SRX130914 | Human* | vaginal* |  |
| PRJNA52107 | SAMN01057066 | SRX155504 | Human* | vaginal* |  |
| PRJNA52105 | SAMN01057067 | SRX155505 | Human* | vaginal* |  |
| PRJNA52107 | SAMN01057066 | SRX155863 | Human* | vaginal* |  |
| PRJNA52105 | SAMN01057067 | SRX155875 | Human* | vaginal* |  |
| PRJNA379934 | SAMN06624125 | SRX2660270 | Human | eye |  |
| PRJNA222257 | SAMN02369387 | SRX456245 | Human | eye |  |
| PRJNA231221 | SAMN11056458 | SRX5949263 | Human | vaginal | (98) |
| PRJNA547620 | SAMN11973370 | SRX5986001 | Human | vaginal | (99) |
| PRJNA547620 | SAMN11973369 | SRX5986002 | Human | vaginal | (99) |
| PRJNA547620 | SAMN11973371 | SRX5986003 | Human | vaginal | (99) |
| PRJNA557339 | SAMN12395213 | SRX6613945 | Human | vaginal | (100) |
| PRJNA563077 | SAMN12667791 | SRX6959881 | Human | gut | (96) |
| PRJNA563077 | SAMN12667801 | SRX6959883 | Chicken | gut | (96) |
| PRJNA563077 | SAMN12667803 | SRX6959885 | Human | gut | (96) |
| PRJNA563077 | SAMN12667804 | SRX6959886 | Turkey | gut | (96) |
| PRJNA563077 | SAMN12667805 | SRX6959887 | Human | eye | (96) |
| PRJNA563077 | SAMN12667793 | SRX6959888 | Chicken | gut | (96) |
| PRJNA563077 | SAMN12667794 | SRX6959889 | Chicken | gut | (96) |
| PRJNA563077 | SAMN12667795 | SRX6959890 | Chicken | gut | (96) |
| PRJNA563077 | SAMN12667796 | SRX6959891 | Chicken | gut | (96) |
| PRJNA563077 | SAMN12667797 | SRX6959892 | Chicken | gut | (96) |
| PRJNA563077 | SAMN12667798 | SRX6959893 | Chicken | gut | (96) |
| PRJNA563077 | SAMN12667799 | SRX6959894 | Chicken | gut | (96) |
| PRJNA563077 | SAMN12667800 | SRX6959895 | Chicken | gut | (96) |
| PRJNA531669 | SAMN11372136 | GCF_009769205 | Chicken | gut | (101) |
| PRJNA231221 | SAMN11056458 | GCF_009730275 | Human | vaginal | (98) |
| PRJNA431864 | SAMN08409124 | GCF_003971565 | Human | vaginal | (102) |
| PRJNA499123 | SAMN10343598 | GCF_003795065 | Human | vaginal | (103) |

Lastly, we looked at patterns of antibiotic resistance across the genus using a table, generated by the *summary* BaT, of resistance genes and loci called by AMRFinder+ (104). Only 79 out of 1,496 *Lactobacillus* samples defined by GTDB-Tk (46) were found to have predicted resistance using AMRFinder+. The most common resistance categories were tetracyclines (67 samples), followed by macrolides, lincosamides and aminoglycosides (16, 15 and 11, respectively). Species with the highest proportion of resistance included *L. amylovorus* (12/14 tetracycline resistant) and *L. crispatus* (10/42 tetracycline resistant). Only 3 genomes of *L. amylophilus* were included in the study but each contained matches to genes for macrolide, lincosamide and tetracycline resistance. The linking thread between these species is they are each commonly isolated from agricultural animals. The high proportion of *L. crispatus* samples isolated from chickens that were tetracycline resistant has been previously observed (105, 106) (**Figure 3**).

A recent analysis of 184 *Lactobacillus* type strain genomes by Campedelli et al (107) found a higher percentage of type strains with aminoglycoside (20/184), tetracycline (18/184), erythromycin (6/184) and clindamycin (60/184) resistance. Forty-two of the type strains had chloramphenicol resistance genes, where here, the AMRFinder+ returned only 1/1,467. These differences probably reflect a combination of the different sampling bias of the studies, and the Campedelli et al strategy of using a relaxed threshold for hits to maximize sensitivity (blastp matches against the CARD database with acid sequence identity of 30% and query coverage of 70% (107)). Resistance is probably undercalled in both methods because of a lack of well-characterized resistance loci from the *Lactobacillus* genus to use as comparison.

## Conclusions

Bactopia is a flexible workflow for bacterial genomics. It can be run on a laptop for a single bacterial sample, but critically, the underlying Nextflow framework allows it to make efficient use of large clusters and cloud-compute environments to process the many thousands of genomes that are currently being generated. For users that are not familiar with bacterial genomic tools and/or who require a standardized pipeline, Bactopia is a “one-stop-shop” that can be easily deployed using conda, Docker and Singularity containers. For researchers with particular interest in individual species or genera, BaDs can be highly customized with taxon-specific databases.

The current version of Bactopia does not support long-read data, but this is planned for future implementation. We also plan to implement more comparative analyses in the form of additional BaTs. With a framework set in place for developing BaTs, it should be possible to make a toolshed of workflows that not only can be used for all bacteria but are also customized for annotating genes and loci specific for particular species.

## Acknowledgements

We would like to thank Torsten Seemann, Oliver Schwengers, Narciso Quijada, Michelle Su, Michelle Wright, Matt Plumb, Sean Wang, Ahmed Babiker and Monica Farley for their helpful suggestions and feedback. We would also like to acknowledge our gratitude to the many scientists and their funders who provided genome sequences to the public domain, ENA and SRA for storing and organizing the data, and the authors of the open source software tools and datasets used in this work.

Support for this project came from the award CDC U50CK000485 Emerging Infection Program (EIP) PPHF/ACA: Enhancing Epidemiology and Laboratory Capacity” funding from the Emory Public Health Bioinformatics Fellowship.

## Availability of data

Raw Illumina sequences used in this study were acquired from Experiment accessions under the BioProject accessions PRJDB1101, PRJDB1726, PRJDB4156, PRJDB4955, PRJDB5065, PRJDB5206, PRJDB6480, PRJDB6495, PRJEB10572, PRJEB11980, PRJEB14693, PRJEB18589, PRJEB19875, PRJEB21025, PRJEB21680, PRJEB22112, PRJEB22252, PRJEB23845, PRJEB24689, PRJEB24698, PRJEB24699, PRJEB24700, PRJEB24701, PRJEB24713, PRJEB24715, PRJEB25194, PRJEB2631, PRJEB26638, PRJEB2824, PRJEB29398, PRJEB29504, PRJEB2977, PRJEB3012, PRJEB3060, PRJEB31213, PRJEB31289, PRJEB31301, PRJEB31307, PRJEB5094, PRJEB8104, PRJEB8721, PRJEB9718, PRJNA165565, PRJNA176000, PRJNA176001, PRJNA183044, PRJNA184888, PRJNA185359, PRJNA185406, PRJNA185584, PRJNA185632, PRJNA185633, PRJNA188920, PRJNA188921, PRJNA196697, PRJNA212644, PRJNA217366, PRJNA218804, PRJNA219157, PRJNA222257, PRJNA224116, PRJNA227106, PRJNA227335, PRJNA231221, PRJNA234998, PRJNA235015, PRJNA235017, PRJNA247439, PRJNA247440, PRJNA247441, PRJNA247442, PRJNA247443, PRJNA247444, PRJNA247445, PRJNA247446, PRJNA247452, PRJNA254854, PRJNA255080, PRJNA257137, PRJNA257138, PRJNA257139, PRJNA257141, PRJNA257142, PRJNA257182, PRJNA257185, PRJNA257853, PRJNA257876, PRJNA258355, PRJNA258500, PRJNA267549, PRJNA269805, PRJNA269831, PRJNA269832, PRJNA269860, PRJNA269905, PRJNA270961, PRJNA270962, PRJNA270963, PRJNA270964, PRJNA270965, PRJNA270966, PRJNA270967, PRJNA270968, PRJNA270969, PRJNA270970, PRJNA270972, PRJNA270973, PRJNA270974, PRJNA272101, PRJNA272102, PRJNA283920, PRJNA289613, PRJNA29003, PRJNA291681, PRJNA296228, PRJNA296248, PRJNA296274, PRJNA296298, PRJNA296309, PRJNA296751, PRJNA296754, PRJNA298448, PRJNA299992, PRJNA300015, PRJNA300023, PRJNA300088, PRJNA300119, PRJNA300123, PRJNA300179, PRJNA302242, PRJNA303235, PRJNA303236, PRJNA305242, PRJNA306257, PRJNA309616, PRJNA312743, PRJNA315676, PRJNA316969, PRJNA322958, PRJNA322959, PRJNA322960, PRJNA322961, PRJNA336518, PRJNA342061, PRJNA342757, PRJNA347617, PRJNA348789, PRJNA376205, PRJNA377666, PRJNA379934, PRJNA381357, PRJNA382771, PRJNA388578, PRJNA392822, PRJNA397632, PRJNA400793, PRJNA434600, PRJNA436228, PRJNA474823, PRJNA474907, PRJNA476494, PRJNA477598, PRJNA481120, PRJNA484967, PRJNA492883, PRJNA493554, PRJNA496358, PRJNA50051, PRJNA50053, PRJNA50055, PRJNA50057, PRJNA50059, PRJNA50061, PRJNA50063, PRJNA50067, PRJNA50115, PRJNA50117, PRJNA50125, PRJNA50133, PRJNA50135, PRJNA50137, PRJNA50139, PRJNA50141, PRJNA50159, PRJNA50161, PRJNA50163, PRJNA50165, PRJNA50167, PRJNA50169, PRJNA50173, PRJNA504605, PRJNA504734, PRJNA505088, PRJNA52105, PRJNA52107, PRJNA52121, PRJNA525939, PRJNA530250, PRJNA533291, PRJNA533837, PRJNA542049, PRJNA542050, PRJNA542054, PRJNA543187, PRJNA544527, PRJNA547620, PRJNA552757, PRJNA554696, PRJNA554698, PRJNA557339, PRJNA562050, PRJNA563077, PRJNA573690, PRJNA577465, PRJNA578299, PRJNA68459, and PRJNA84.

## Links

- Website/Docs - https://bactopia.github.io/
- Github - https://www.github.com/bactopia/bactopia/
- Zenodo Snapshot - https://doi.org/10.5281/zenodo.3691353
- Bioconda - https://bioconda.github.io/recipes/bactopia/README.html
- Containers

○ Docker - https://cloud.docker.com/u/bactopia/
○ Singularity - https://cloud.sylabs.io/library/rpetit3/bactopia

## Supplementary Tables

**Supplementary Table 1.**
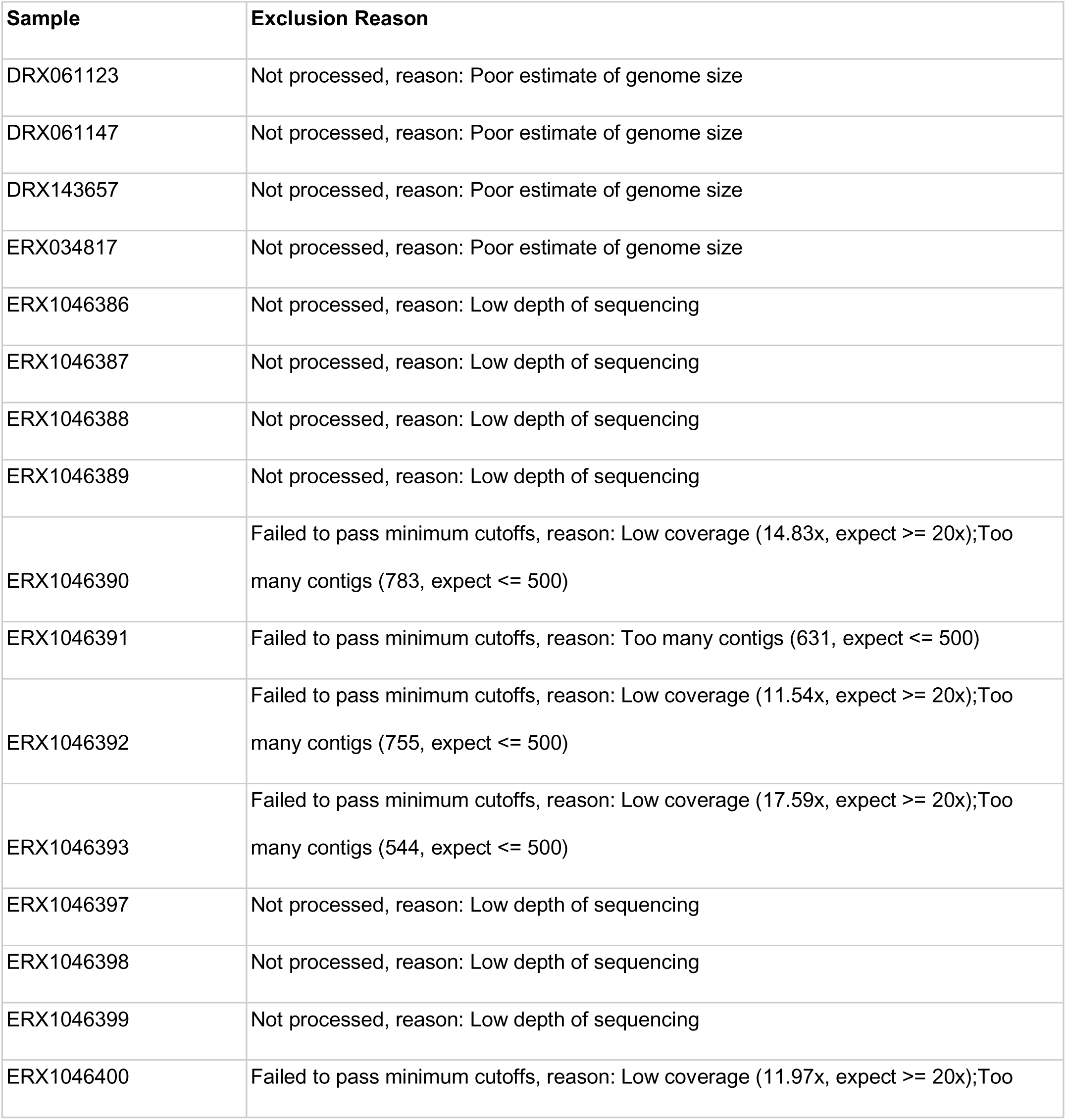
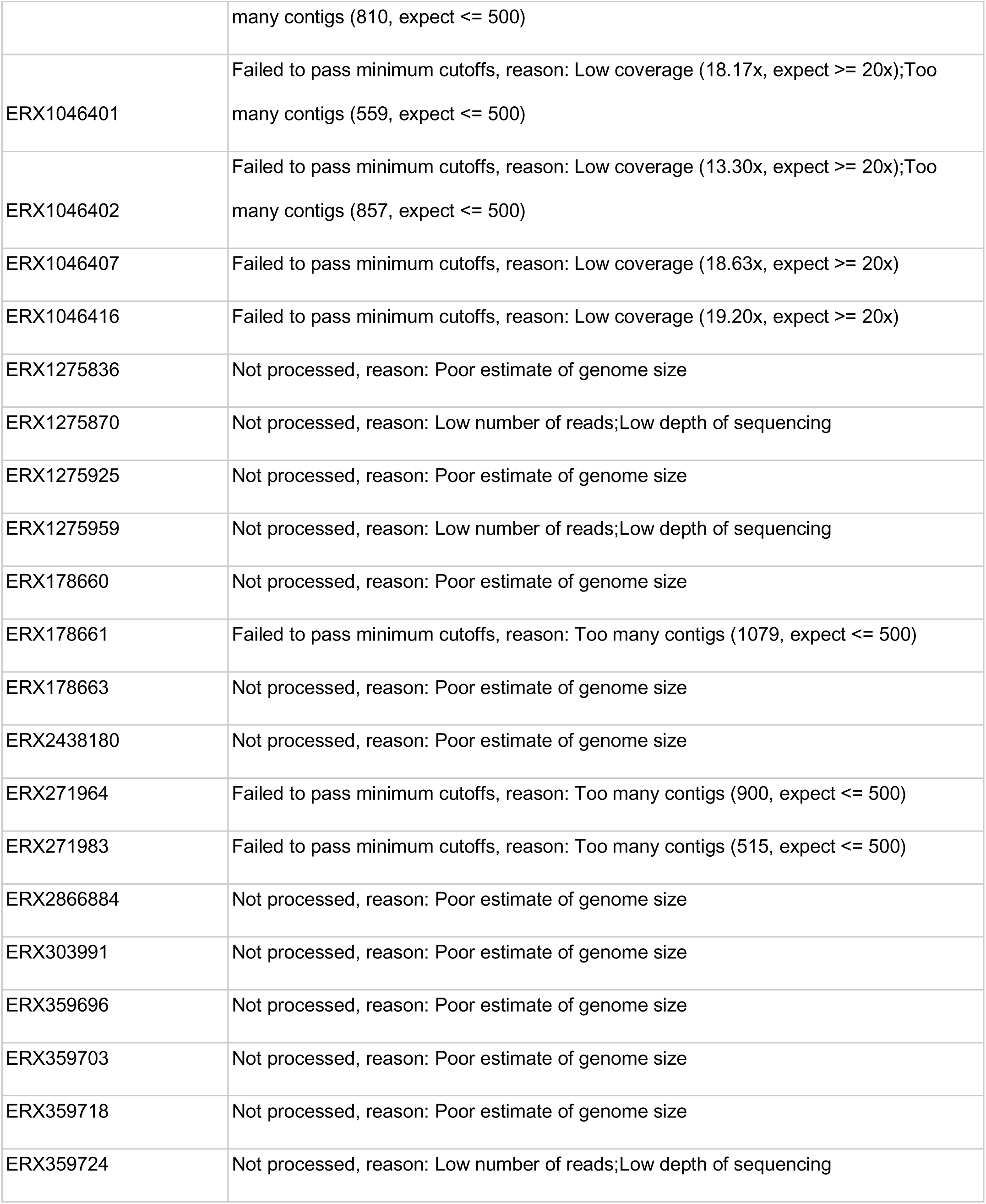
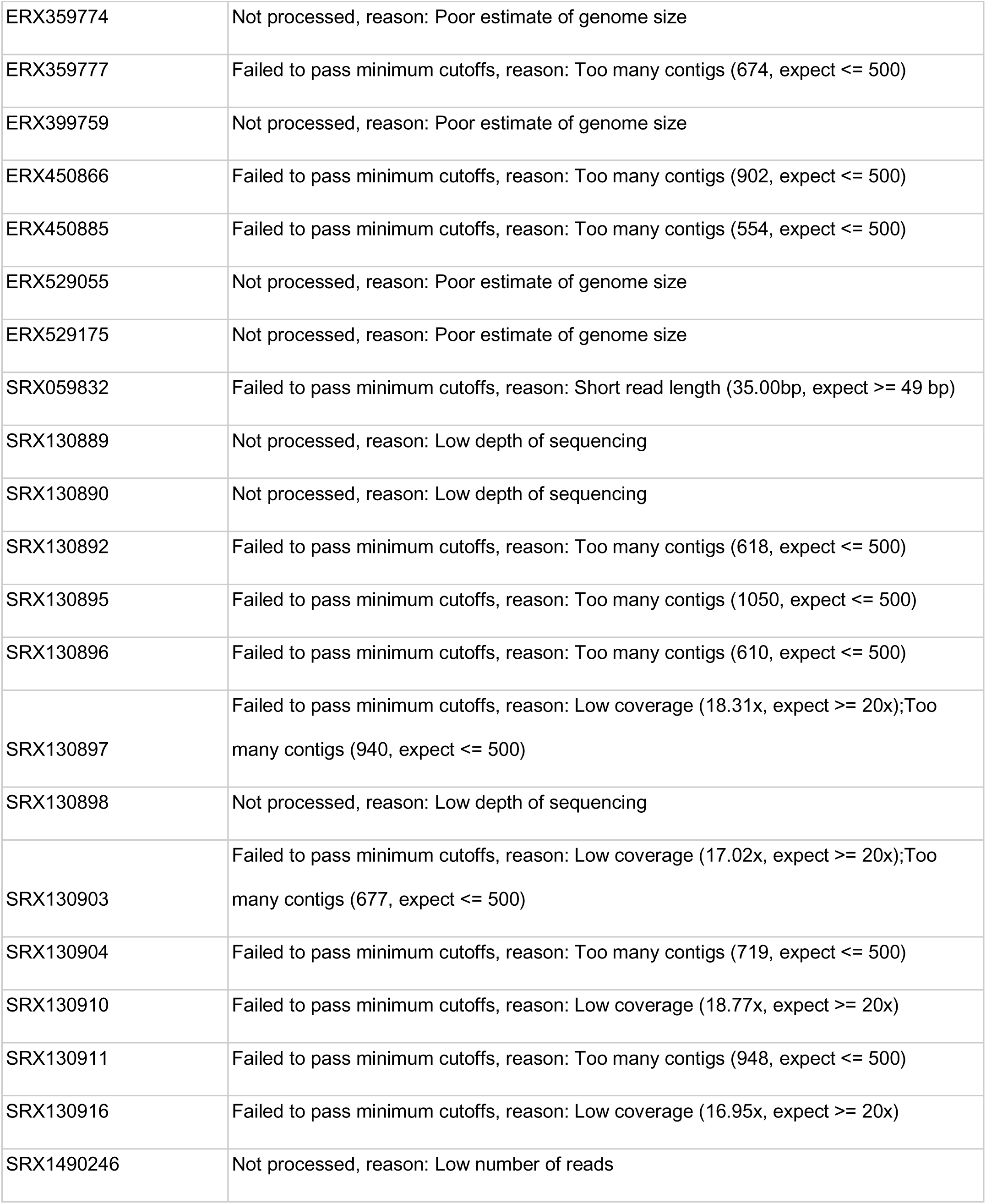
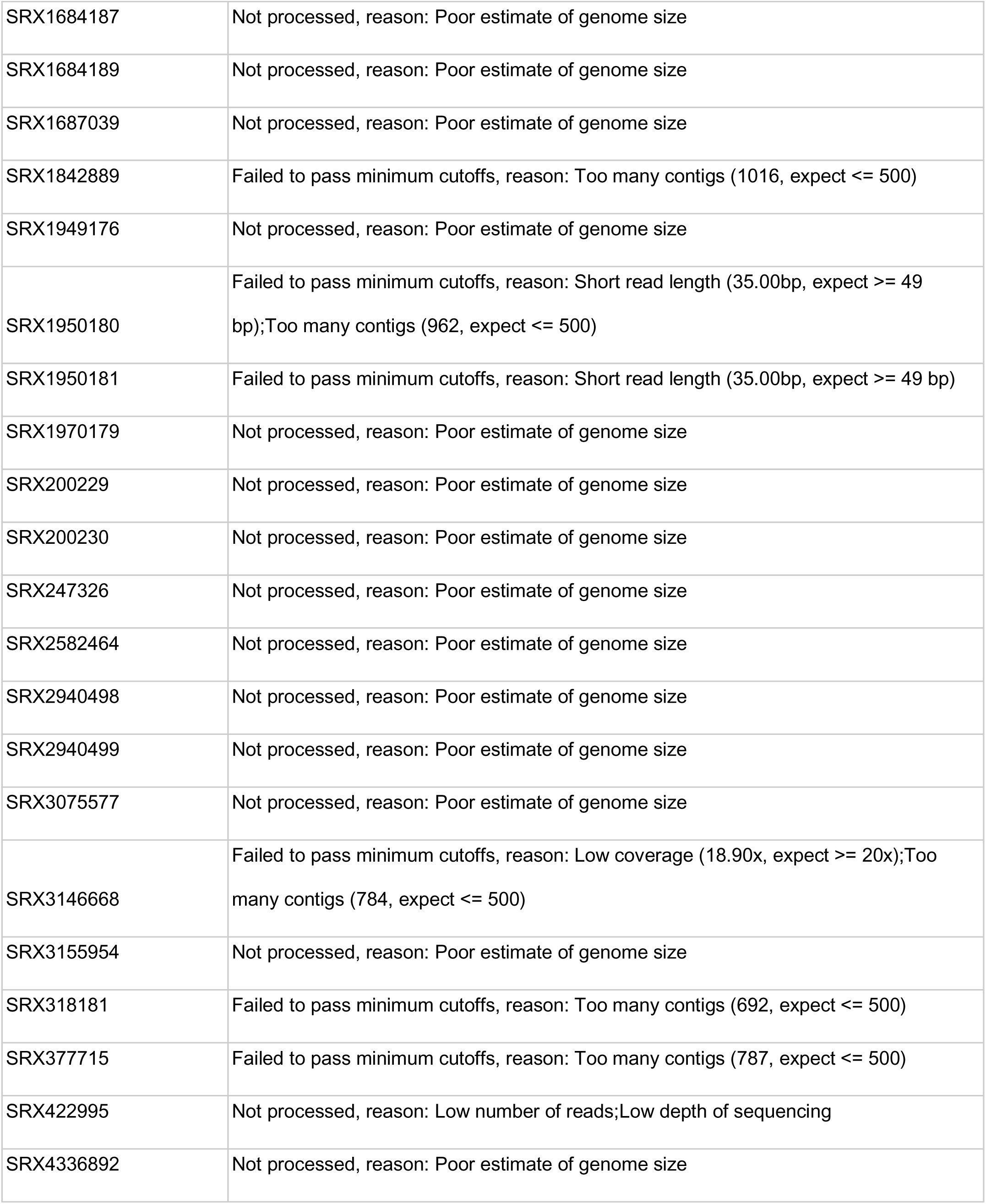
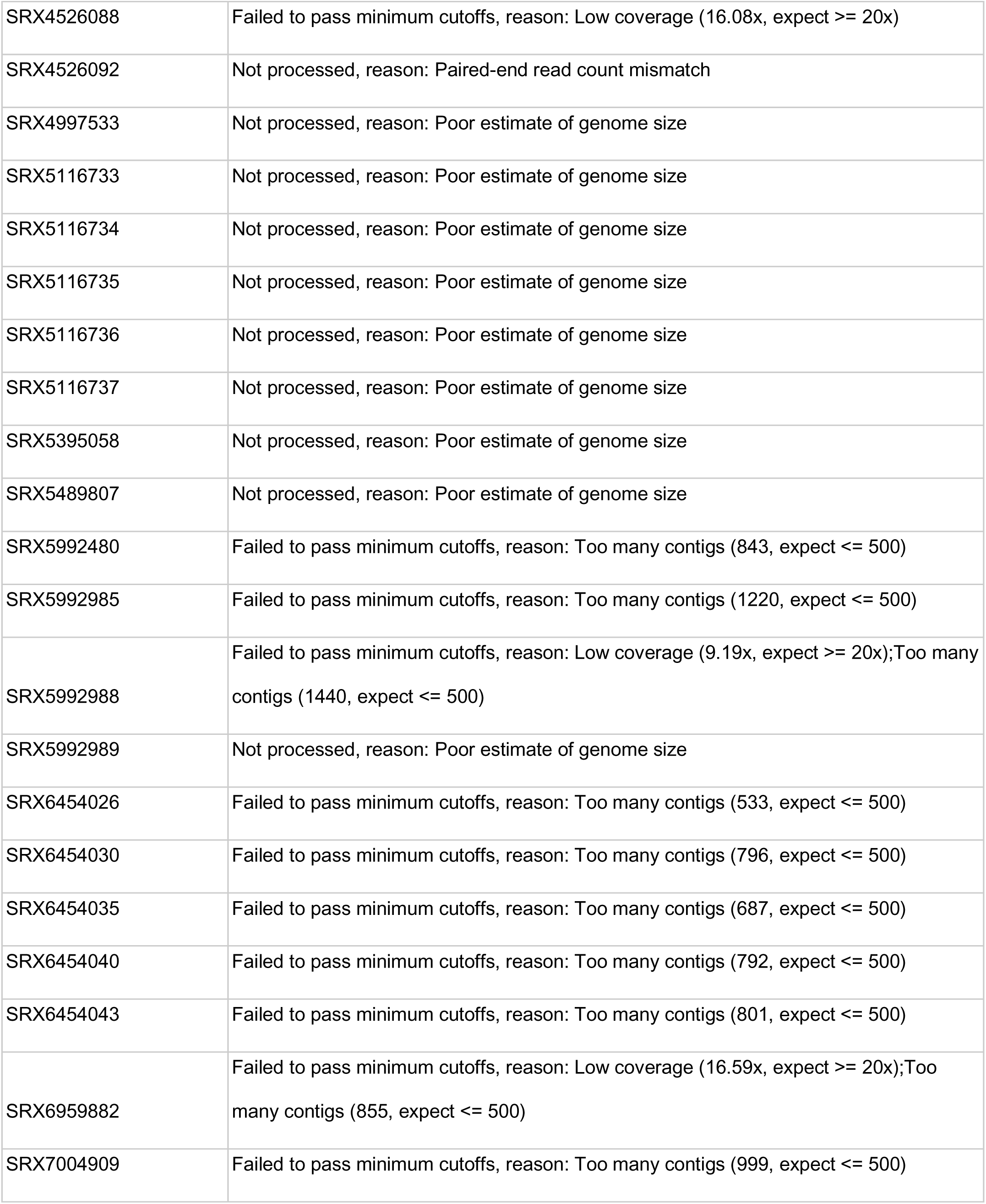
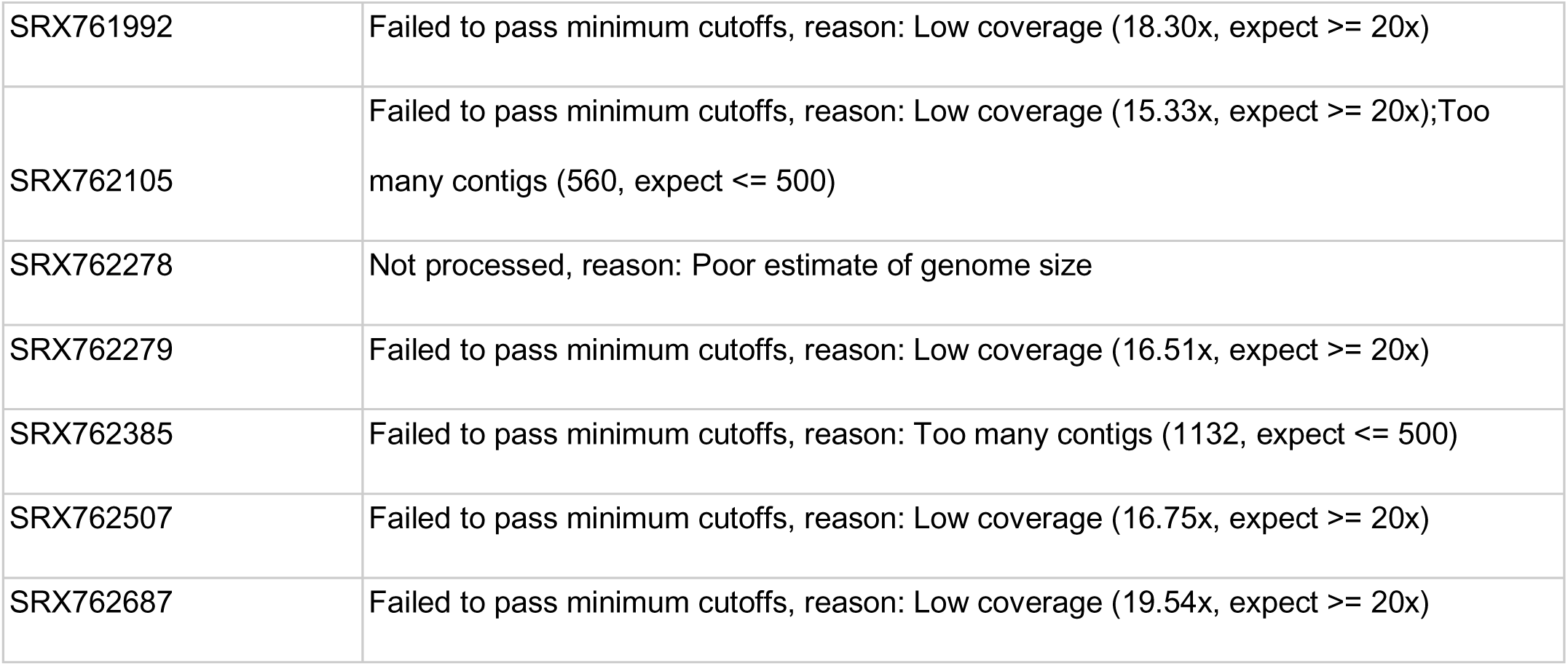
*Lactobacillus* excluded from analysis.

**Supplementary Table 2.**
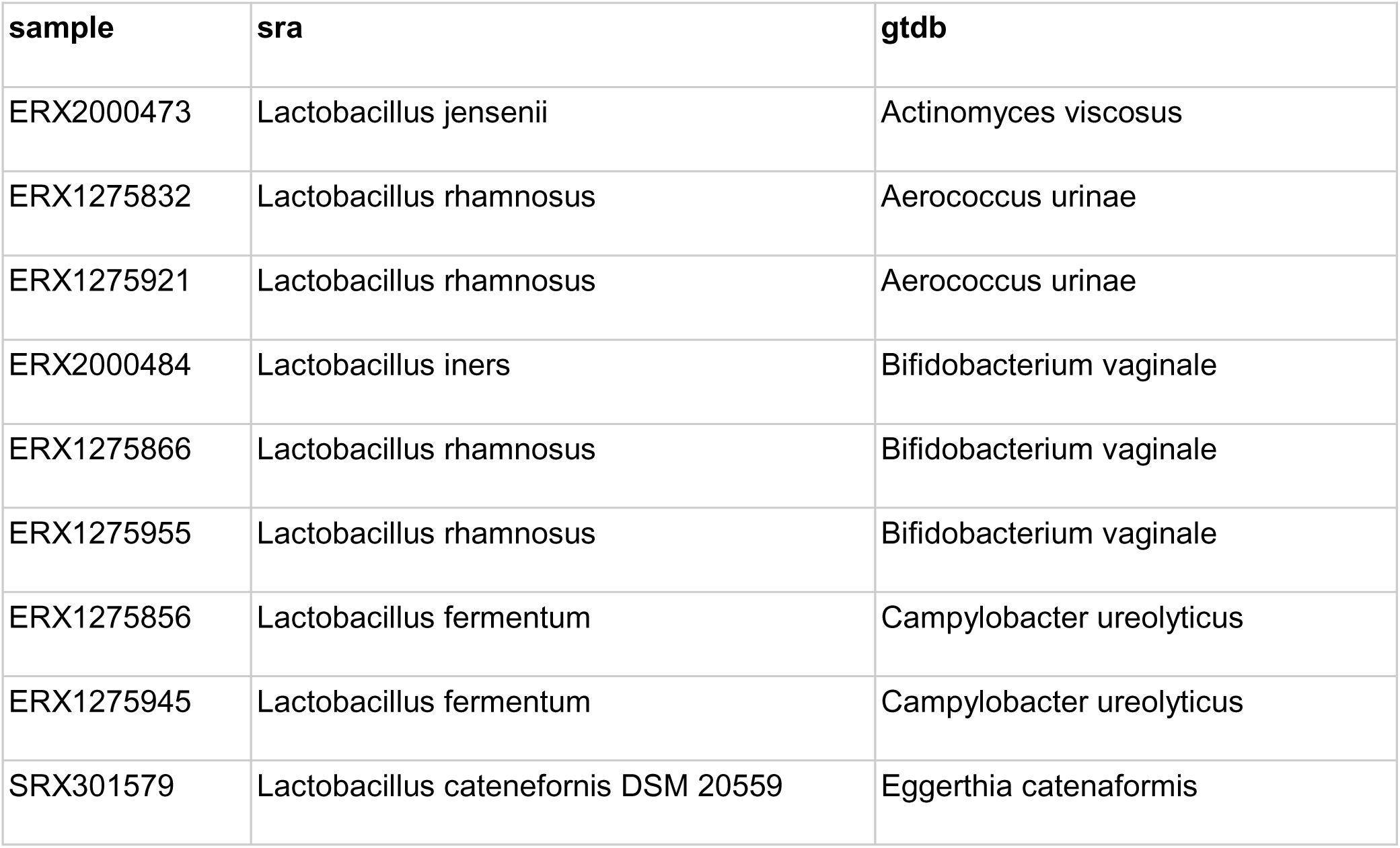
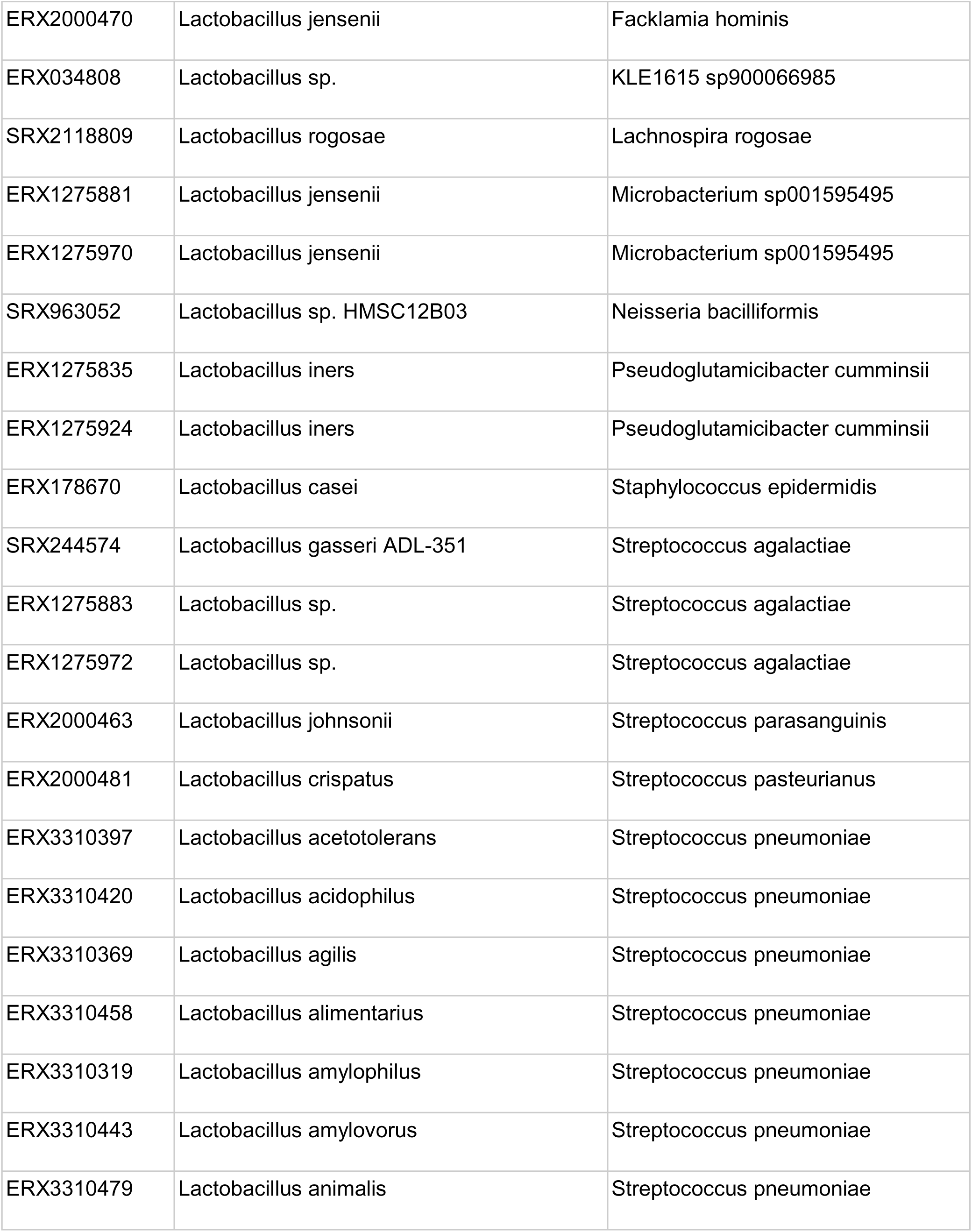
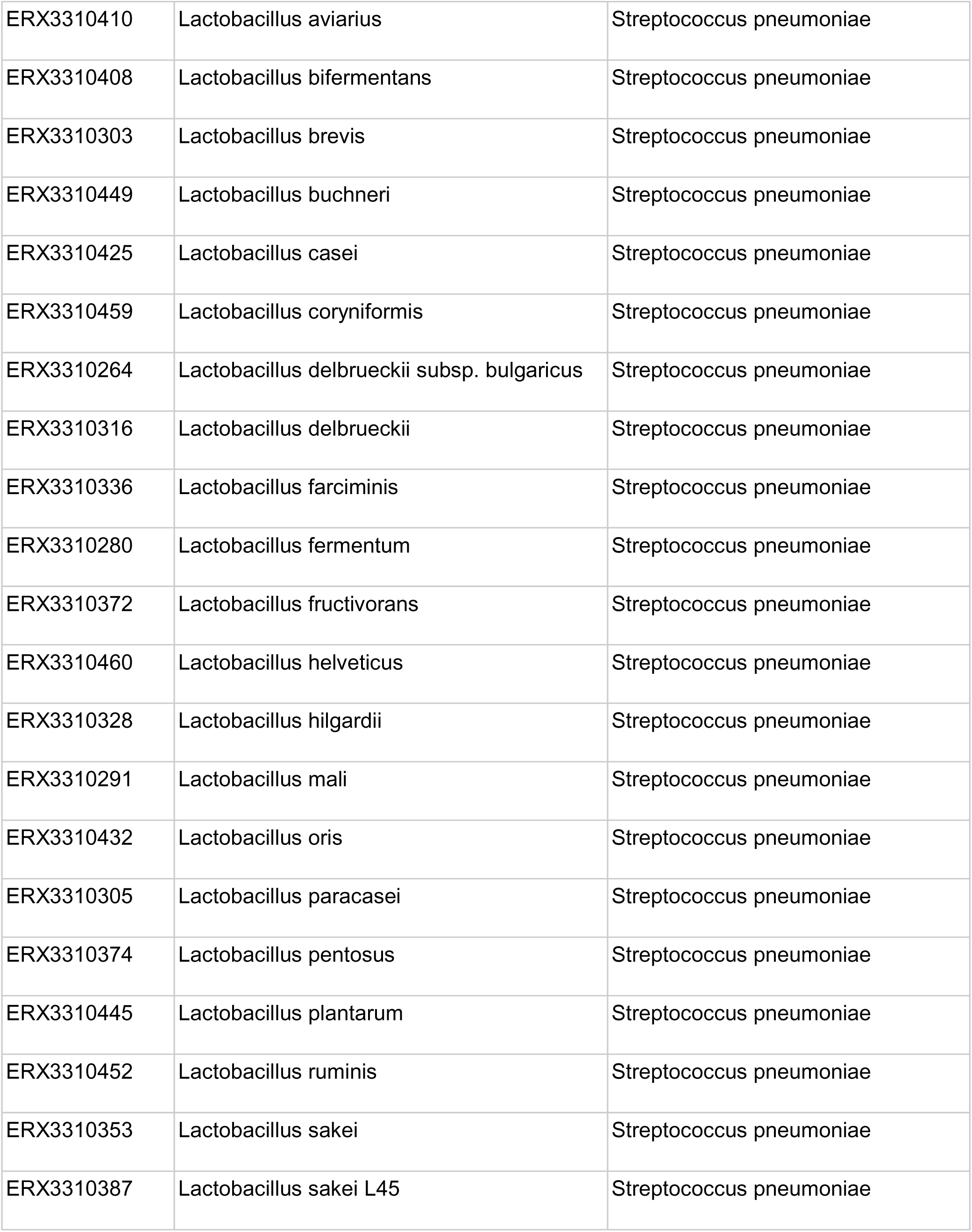
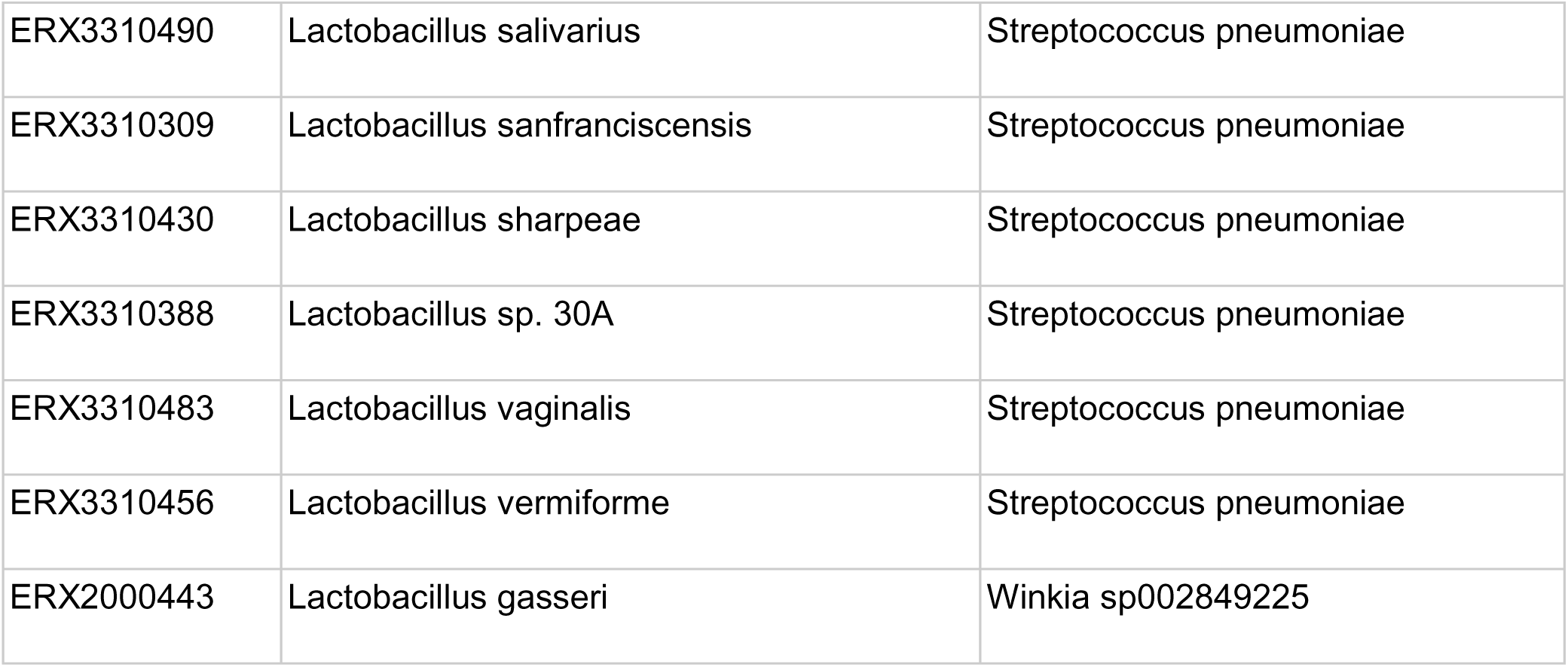
Samples with a non-Lactobacillus taxonomic classification.

## Supplementary Figures

**Supplementary Figure 1.**
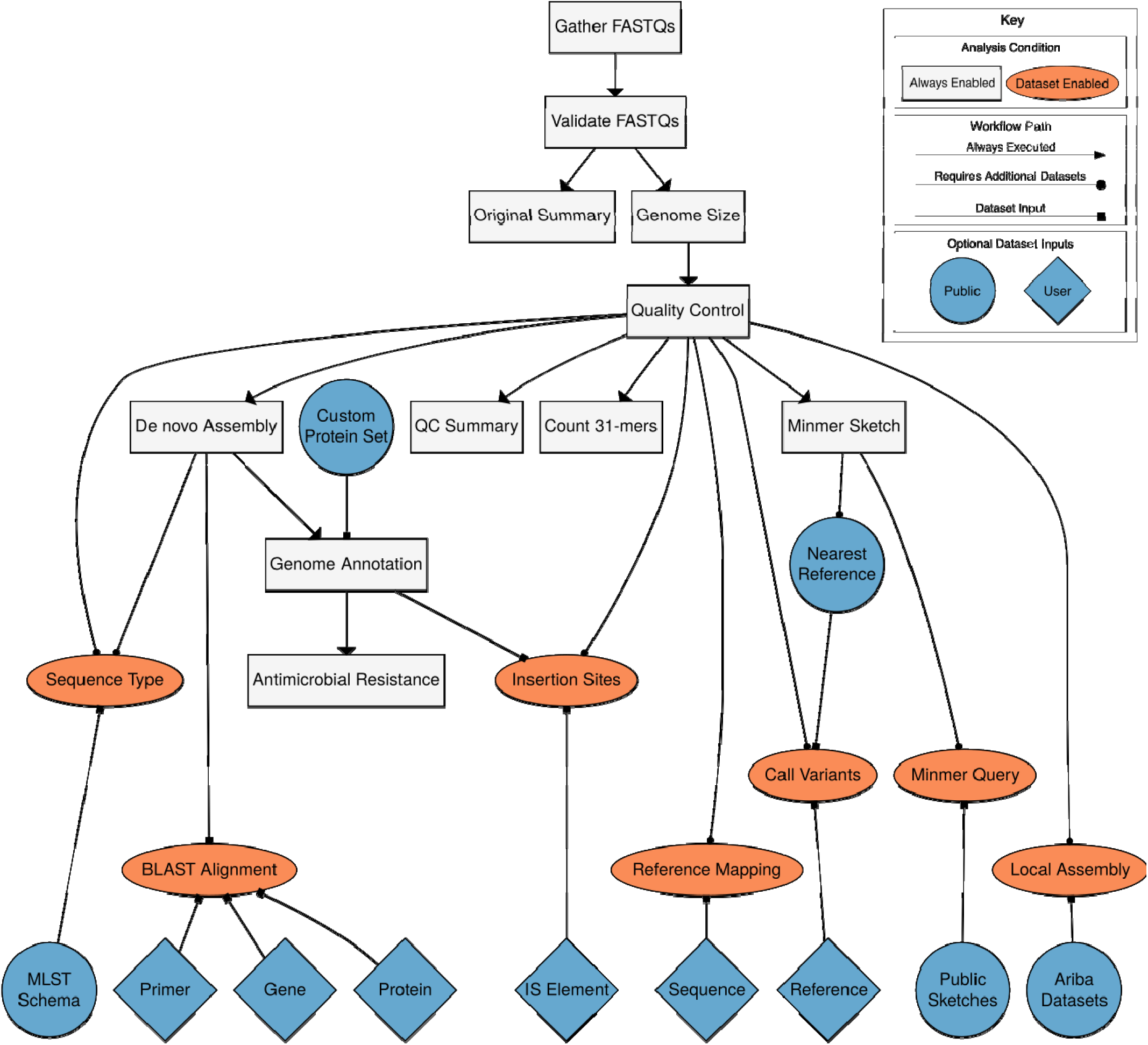
Bactopia Analysis Pipeline Workflow.

**Supplementary Figure 2.**
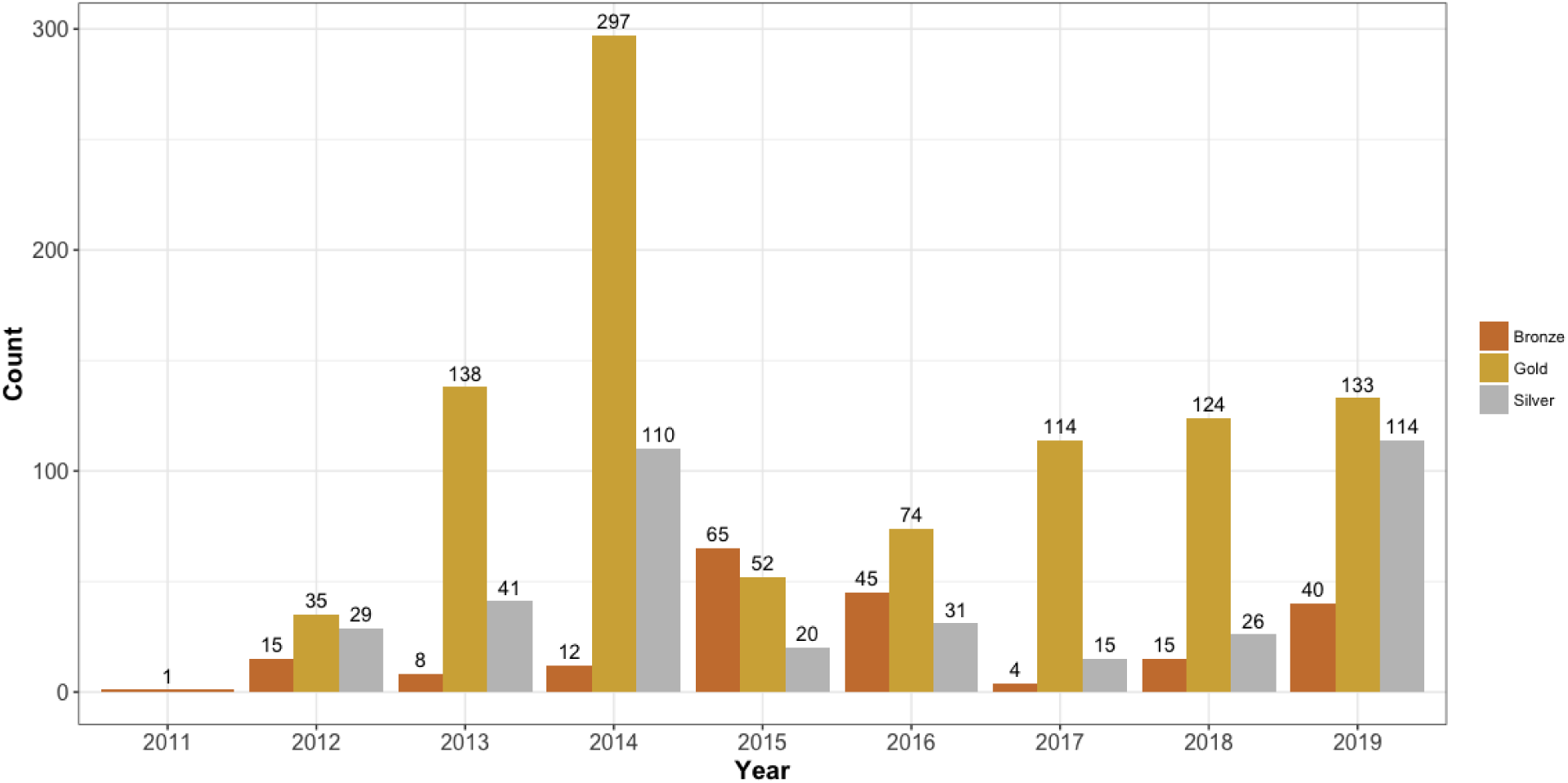
Sequencing quality ranks per year 2011–2019 of Lactobacillus genome projects. Genome projects were grouped into three increasing quality ranks: Bronze, Silver and Gold. The rank was based on coverage, read length, per-read quality and total assembled contigs. The highest rank, Gold, represented 62% (n=967) of the available Lactobacillus genome projects. The remaining genomes were 25% Silver (n=386) and 13% Bronze (n=205). Between the years of 2011 and 2019, Gold rank samples consistently outnumbered Silver and Bronze except for 2011 and 2015. However it is likely the total number of Gold rank samples is underrepresented due coverage reduction being based on the estimated genome size.

**Supplementary Figure 3.**
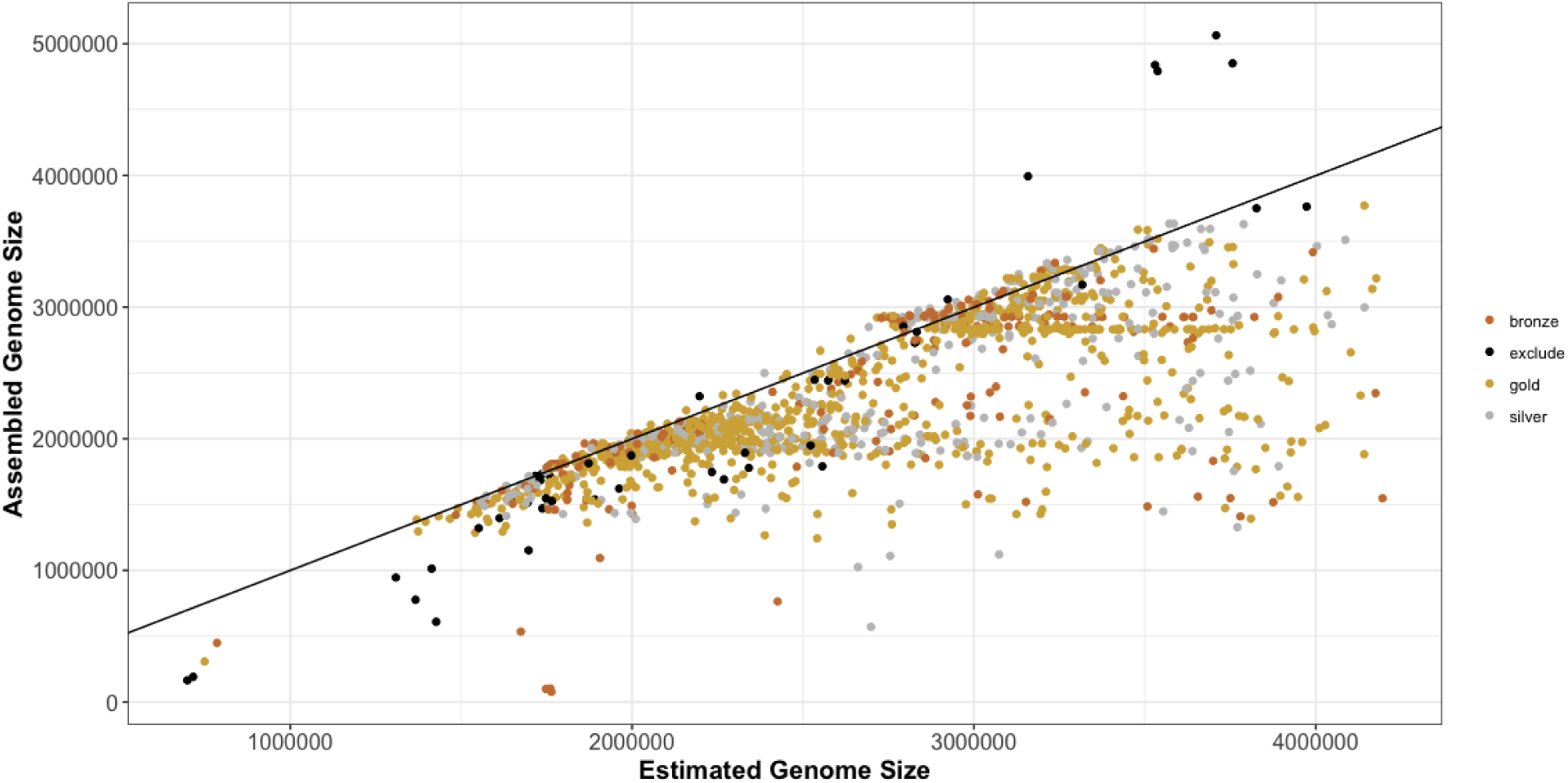
Comparison of estimated genome size and assembled genome size. The assembled genome size (Y-axis) and estimated genome size (X-axis) was plotted for each sample. The color of the dot is determined by the rank of the sample. The solid black line represents a 1:1 ratio between the assembled genome size and the estimated genome size. The genome size was estimated for each sample by Mash (7) using the raw sequences.

**Supplementary Figure 4.**
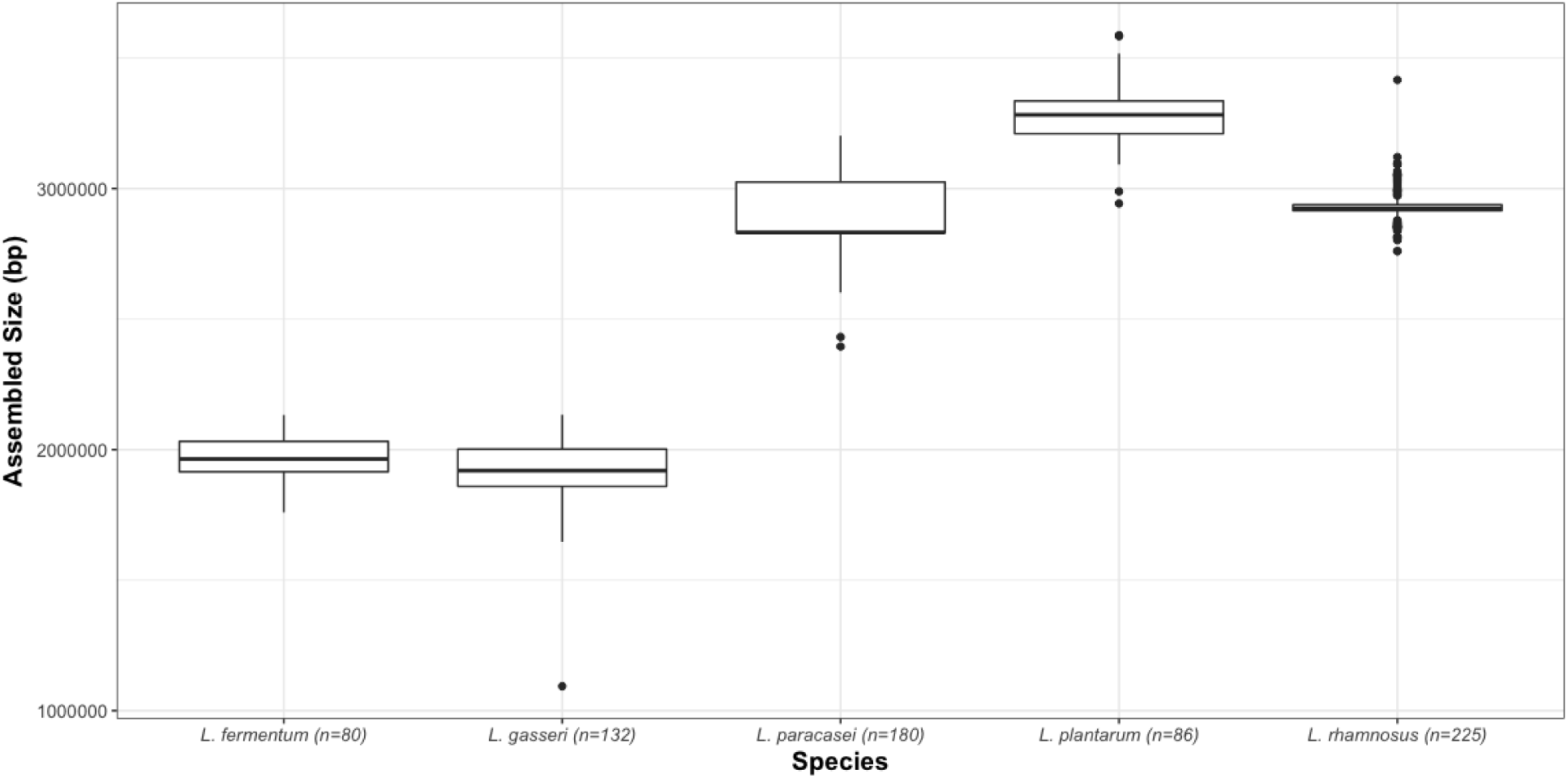
Assembled genome sizes are consistent within species.

## Supplementary Data

Supplementary Data 1 - Results returned after querying ENA for Lactobacillus

Supplementary Data 2 - SRA/ENA Experiment accessions processed by Bactopia

Supplementary Data 3 - Nexflow runtime report for *Lactobacillus* genomes processed by Bactopia

