## Supplemental Data 1 for "Bactopia: a flexible pipeline for complete analysis of bacterial genomes": supplementary-data-3-nextflow-report.html

[cranky\_torricelli] Nextflow Workflow Report


Nextflow Report


- Summary
- Resources
- Tasks

[cranky\_torricelli]

### Nextflow workflow report

#### `[cranky_torricelli]`

Workflow execution completed successfully!

Run times
:   05-Nov-2019 08:06:30 - 07-Nov-2019 20:21:36
    (duration: **2d 12h 15m 6s**)

31295 succeeded

0 cached

0 ignored

30 failed

Nextflow command
:   ```
    nextflow /home/rpetit3/repos/bactopia/main.nf --accessions ../lactobacillus-accessions.txt --datasets /home/rpetit3/datasets --species lactobacillus --coverage 100 --cpus 4 -profile slurm --min_genome_size 1000000 --max_genome_size 4200000
    ```

CPU-Hours
:   `3'052.4 (0% failed)`

Launch directory
:   `/home/rpetit3/projects/lactobacillus/bactopia`

Work directory
:   `/home/rpetit3/projects/lactobacillus/bactopia/work`

Project directory
:   `/home/rpetit3/repos/bactopia`

Script name
:   `main.nf`

Script ID
:   `b763323ed40fb19eda2aff0c1a481f11`

Workflow session
:   `90a3fb3c-b891-4c5a-a6b5-5c6159277170`

Workflow profile
:   slurm

Nextflow version
:   version 19.10.0, build 5170 (21-10-2019 15:07 UTC)

## Resource Usage

These plots give an overview of the distribution of resource usage for each process.

#### CPU

- Raw Usage
- % Allocated

#### Memory

- Physical (RAM)
- Virtual (RAM + Disk swap)
- % RAM Allocated

#### Job Duration

- Raw Usage
- % Allocated

#### I/O

- Read
- Write

## Tasks

This table shows information about each task in the workflow. Use the search box on the right
to filter rows for specific values. Clicking headers will sort the table by that value and
scrolling side to side will reveal more columns.

Values shown as:

Human readable
Raw values

(tasks table omitted because the dataset is too big)

Generated by Nextflow, version 19.10.0
